# Structural variation at *lhcb6* underlies genetic variation in photosystem II maximum quantum efficiency in maize

**DOI:** 10.1101/2025.09.16.676272

**Authors:** Sebastian Urzinger, Lukas Würstl, Viktoriya Avramova, Claude Urbany, Daniela Scheuermann, Thomas Presterl, Stefan Reuscher, Manfred Mayer, Sarah Brajkovic, Milena Ouzunova, Bernardo Ordas, Peter Westhoff, Chris-Carolin Schön

## Abstract

Targeted utilization of native genetic diversity can expand the genetic basis of traits that exhibit limited genetic variation in elite breeding material and can enhance our understanding of the genotype-phenotype relationships associated with complex traits in crops. In a genome-wide association study in a European maize landrace we identified quantitative trait loci (QTL) that affected the maximum quantum efficiency of photosystem II (F_v_/F_m_) in field experiments. In a forward genetic approach, we focused on a QTL on chromosome 10, explaining a large proportion of the genetic variance for F_v_/F_m_ in growth stages V4 (35%) and V6 (47%), for genetic dissection and candidate gene discovery. Integrating molecular and physiological information we show that allelic variation at the gene encoding LIGHT HARVESTING CHLOROPHYLL A/B BINDING PROTEIN6 (LHCB6), a component of the photosystem II (PSII) light-harvesting complex (LHCII) antenna, underlies the variation in F_v_/F_m_. We demonstrate that the allelic variation results from a hAT transposon insertion at *lhcb6* and is associated with differential accumulation of the LHCII antenna components LHCB6 and LHCB3, leading to differences in non-photochemical quenching (NPQ) and plant biomass accumulation. Based on proteomic analyses we propose candidate genes that compensate the unfavorable effects caused by impaired LHCII antenna assembly. Our work provides novel insights into the function of *lhcb6* in the context of LHCII antenna assembly and demonstrates the value of natural variation in landraces for the understanding and genetic improvement of complex photosynthetic processes.

## Introduction

Crop yields have increased steadily in recent decades, yet future genetic improvement will require broadening the genetic base of elite crop germplasm (McCouch et al., 2013, Swarup et al., 2021). In maize (*Zea mays* L.), multiple genetic bottlenecks—starting with its geographic expansion from its center of origin, followed by the establishment of heterotic groups using a small number of landraces, and subsequent cycles of intense selection—have significantly narrowed the genetic diversity of elite germplasm (Mir et al., 2013, Gouesnard et al., 2017, Reif et al., 2005). Thus, beneficial alleles, that influence (a)biotic stress tolerance, resource efficiency and photosynthetic traits, likely have been lost over time and undesirable alleles might have been fixed through genetic drift or linkage drag (Hartfield and Otto, 2011, Lobell and Tebaldi, 2014, Sharwood et al., 2022). Breakthroughs in genetic engineering have deepened our understanding of plant physiology and raised hopes for the development of improved crops. However, when it comes to quantitative traits, translating molecular advances to field performance often falls short of expectations (Theeuwen et al., 2022, Khaipho-Burch et al., 2023). Alternatively, landraces offer a wealth of untapped allelic variation that can be leveraged to improve agronomically and ecologically significant traits (McCouch et al., 2013, Hölker et al., 2019, Mayer et al., 2020, Liang et al., 2021). Despite their conceptual value, the incorporation of landraces into breeding programs is hindered by their yield gap compared to elite lines (Hallauer and Sears, 1972, Strigens et al., 2013, Hölker et al., 2019) and the risk of linkage drag due to the co-inheritance of undesirable and beneficial alleles (Sood et al., 2014, Böhm et al., 2017). One strategy to overcome these challenges is to extract clearly defined alleles associated with quantitative traits of interest from preselected landraces (Mayer et al., 2020). In a previous study, we reported a successful example for such an approach by demonstrating that favorable allelic variation at the *ndhm1* locus, modulating cyclic electron transport, has pleiotropic positive effects on photosystem II efficiency, cold tolerance and plant growth (Urzinger et al., 2025).

Here, we focus on the target trait maximum quantum efficiency of photosystem II (PSII, F_v_/F_m_). F_v_/F_m_ provides an estimate of how efficient light energy is used for photochemistry. Its physiological interpretation depends on the plant species, genetic background, and environmental conditions (Murchie and Lawson, 2013). In maize, genetic variation in F_v_/F_m_ has been shown to be correlated with multiple traits such as early plant development, chilling tolerance, PSII operating efficiency (ΦPSII), non-photochemical quenching (NPQ) and chlorophyll content (Hölker et al., 2019, Ferguson et al., 2025, Lainé et al., 2023). Thus, F_v_/F_m_ can serve as a proxy for assessing genetic variation related to stress tolerance (Burnett and Kromdijk, 2022), but it is also related to biomass production by modulation of light interception- and conversion-efficiency (Murchie and Lawson, 2013, Long et al., 2015, Leister, 2023, Theeuwen et al., 2022). Previous studies using bi-parental or multi-parent advanced generation intercross (MAGIC) populations have identified only a few quantitative trait loci (QTL) for F_v_/F_m_, each with relatively large effects (Fracheboud et al., 2002, Fracheboud et al., 2004, Hund et al., 2004, Jompuk et al., 2005, Ferguson et al., 2025). Candidate genes have been proposed for these QTL, but the physiological mechanisms underlying the variation of F_v_/F_m_ in maize have remained largely unclear.

In this study we performed a comprehensive analysis of QTL for F_v_/F_m_ in a population of 222 doubled-haploid (DH) lines derived from the maize landrace “Kemater Landmais Gelb” (Kemater). First, we conducted a genome-wide association study (GWAS), to explore the genetic architecture of F_v_/F_m_ variation in the landrace. We then related the genetic variation in F_v_/F_m_ to physiological parameters significantly influencing agronomically important traits. Our analysis led to the discovery of a genomic region on maize chromosome 10 harboring allelic variation at *lhcb6*, a gene encoding a component of the PSII light-harvesting complex, as a key determinant of F_v_/F_m_ variation in Kemater. We subsequently characterized the impact of this allelic variation on transcript levels and the leaf proteome, as well as on the physiological trait NPQ and plant growth. We also investigated the prevalence of specific alleles at this gene in diverse maize inbred lines.

## Results

### Discovery of QTL affecting Fv/Fm in a maize landrace

In the genome-wide association study (GWAS), significant marker trait associations were detected for maximum quantum efficiency of photosystem II (PSII, F_v_/F_m_) on chromosomes 1 (growth stage V6), 2 (V4, V6), 4 (V4), 8 (V4), and 10 (V4, V6; Table 1). The QTL on chromosomes 1, 4 and 8 were detected in one growth stage only, while the QTL on chromosomes 2 and 10 were discovered in V4 and V6. Because the confidence intervals of the QTL on chromosomes 2 and 10 largely overlapped between the two growth stages, each genomic region was considered a single QTL affecting both growth stages in subsequent analyses. Together, the five QTL at the above-mentioned positions explained 50% and 59% of the genetic variance of F_v_/F_m_ in growth stages V4 and V6, respectively (Table 1). Here, we focus on the QTL on chromosome 10 spanning a 3.9 Mb genomic region at the distal end of the chromosome (Figure 1). This QTL had an effect size on F_v_/F_m_ of 0.45 (0.46) standard deviations (SD) and explained 35% (47%) of the genetic variance in V4 (V6). Due to a large effect in both growth stages a bi-parental population was developed for fine mapping and functional characterization of this region.

**Figure 1.**
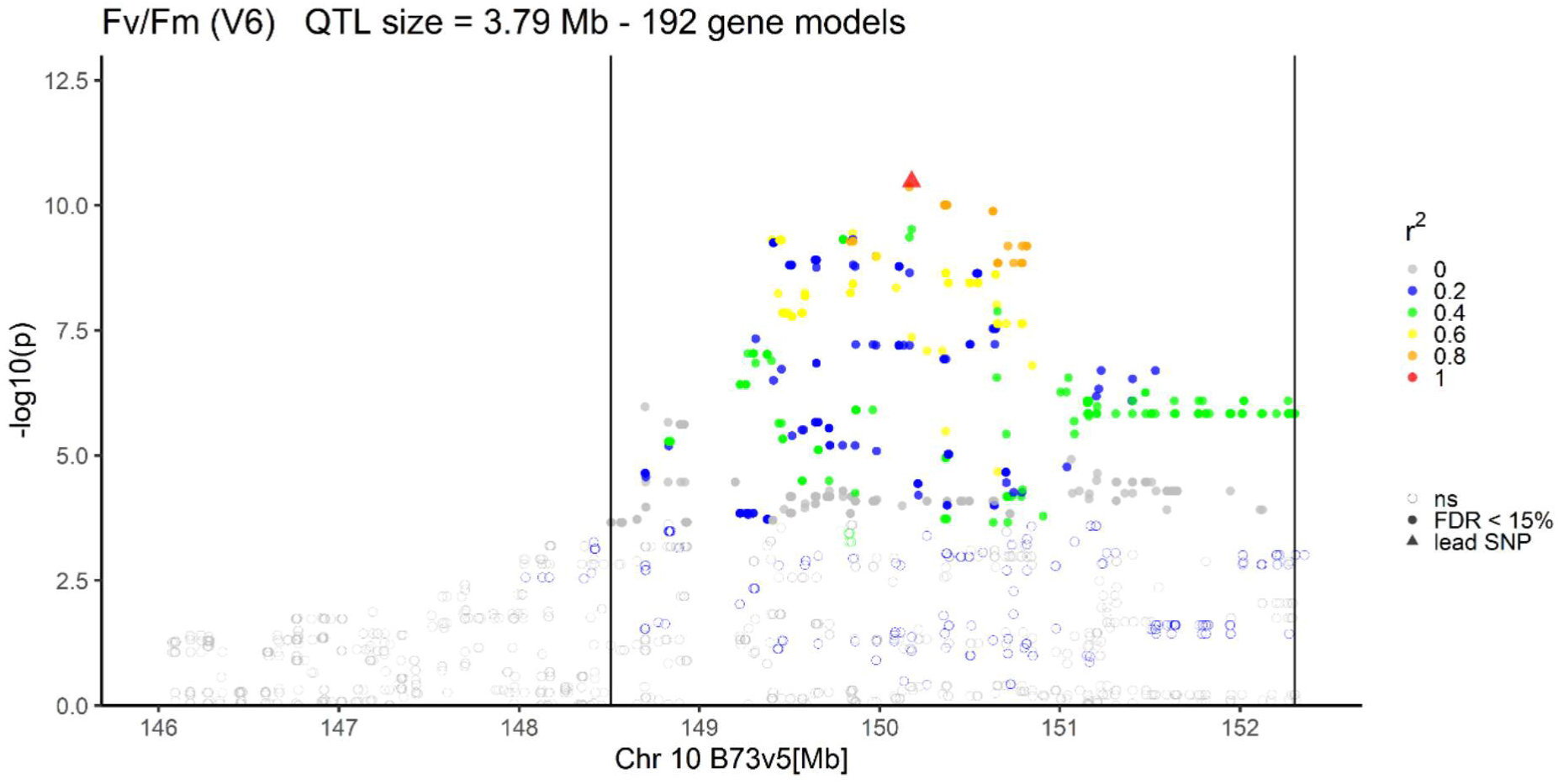
Marker-trait associations from a genome-wide association study (GWAS) for maximum potential quantum yield of photosystem II (F_v_/F_m_) on chromosome 10. Phenotypic data were assessed on 211 DH lines derived from the landrace Kemater during early development in growth stage V6 in field experiments in two locations. The x-axis indicates the physical position of SNPs on chromosome 10 in the reference sequence B73v5. Significance of associations of SNPs with F_v_/F_m_ (false discovery rate: FDR < 15%) is indicated on the y-axis and by symbols: empty circles (no significant association, ns); filled circles (significant association; FDR < 15%); triangle (SNP with highest significance, lead SNP, AX-90599221). Linkage disequilibrium (r^2^) between the lead SNP and SNPs in the target region is color-coded. QTL boundaries are given by vertical lines.

**Table 1.** QTL for maximum potential quantum yield of photosystem II (F_v_/F_m_) in landrace Kemater. The genome-wide association study (GWAS) was conducted using approximately 500,000 SNPs and adjusted entry means of F_v_/F_m_ in growth stages V4 and V6. Genomic positions are provided in B73v5 coordinates, and the number of genes within each QTL region is indicated. Effect SD: Absolute effect size from GEMMA output standardized by the standard deviation of the respective trait. GVE: Genetic variance explained by the respective QTL. Total GVE: Genetic variance explained by a model fitting all detected QTL in a growth stage simultaneously.

| QTL | Growth stage | Chr | Start | End | Genes | Effect SD | GVE | Total GVE |
| --- | --- | --- | --- | --- | --- | --- | --- | --- |
| qV4FvFm2.1 | V4 | 2 | 22392853 | 25329743 | 79 | -0.43 | 7% | 50% |
| qV4FvFm4.1 | V4 | 4 | 244037990 | 244041963 | 1 | -0.32 | 7% |  |
| qV4FvFm8.1 | V4 | 8 | 40669624 | 40670607 | 1 | 1.35 | 5% |  |
| qV4FvFm10.1 | V4 | 10 | 148413861 | 152303594 | 194 | -0.45 | 35% |  |
| qV6FvFm1.1 | V6 | 1 | 160150163 | 163660341 | 25 | 0.33 | 30% | 59% |
| qV6FvFm2.1 | V6 | 2 | 24398414 | 27269195 | 81 | -0.39 | 12% |  |
| qV6FvFm10.1 | V6 | 10 | 148506993 | 152303594 | 192 | -0.46 | 47% |  |

### Genetic dissection of the genomic region on chromosome 10

Two Kemater DH lines, KE0095 and KE0109, were selected from the GWAS population as parents of a bi-parental population (Figure 2A), based on their phenotypic difference in F_v_/F_m_ (Figure S1) and their genomic similarity assessed with the 600k Axiom™ Maize SNP Array (600k Array; Unterseer et al., 2014). The two lines are polymorphic in the target region on chromosome 10 (Chr10: 148,413,861-152,303,534; Figure 1), share the same SNP allele at the other four F_v_/F_m_ QTL (Table 1) and have a similar genomic background with 419,014 out of 493,989 SNPs being monomorphic (Figure S2). Chromosome 10 is nearly identical between KE0095 and KE0109 based on sequence alignment (Figure 2B) and 30,120 out of 32,236 SNPs being monomorphic. Polymorphic SNPs on chromosome 10 were confined to a 22 Mb region (Chr10: 130,130,769–152,357,073), covering the QTL for F_v_/F_m_. Thus, the bi-parental population allowed fine-mapping of the target region (Chr10: 148,413,861-152,303,534) with no interference of other QTL. F_2:3_ lines derived from the KE0095 x KE0109 cross, showing recombination events in the target region, were phenotyped for F_v_/F_m_ at three German field locations Freising (FRS), Roggenstein (ROG) and Einbeck (EIN) in 2024. Entry-mean heritability (h^2^) for F_v_/F_m_ was 0.63 (95% CI: 0.40 – 0.77), indicating a significant genetic component underlying trait expression in the F_2:3_ recombinant lines. The association with F_v_/F_m_ was assessed for 14 markers in the target region, of which eight were significantly associated with F_v_/F_m_ (false discovery rate: FDR < 1%; Figure 3A). The strongest association with F_v_/F_m_ was observed for the most significant SNP from the GWAS analysis (AX-90599221; lead SNP; Table S1). The KE0095 allele of the lead SNP had a trait increasing effect on F_v_/F_m_ compared to the KE0109 allele, as expected from trait expression of the parents (Figure S1). This marker explained 56% of the phenotypic and 98% of the genetic variance in the 52 F_2:3_ lines, suggesting that the underlying gene is the major determinant of F_v_/F_m_ segregation in this bi- parental population. The strength of the association with F_v_/F_m_ diminished by several orders of magnitude up- and downstream of the lead SNP (Figure 3A). Recombination events were observed between the lead SNP and neighboring SNPs (AX-91369070, AX-91196092; maximum r^2^=0.71; Figure 3B), allowing fine-mapping of the region. The fine-mapping narrowed the genomic region associated with F_v_/F_m_ to a 154 kb fragment between the markers AX-91369070 and AX-91196092 (Chr10:150,109,398-150,262,989; blue box in Figure 3A, Figure 3B, Table S1).

**Figure 2.**
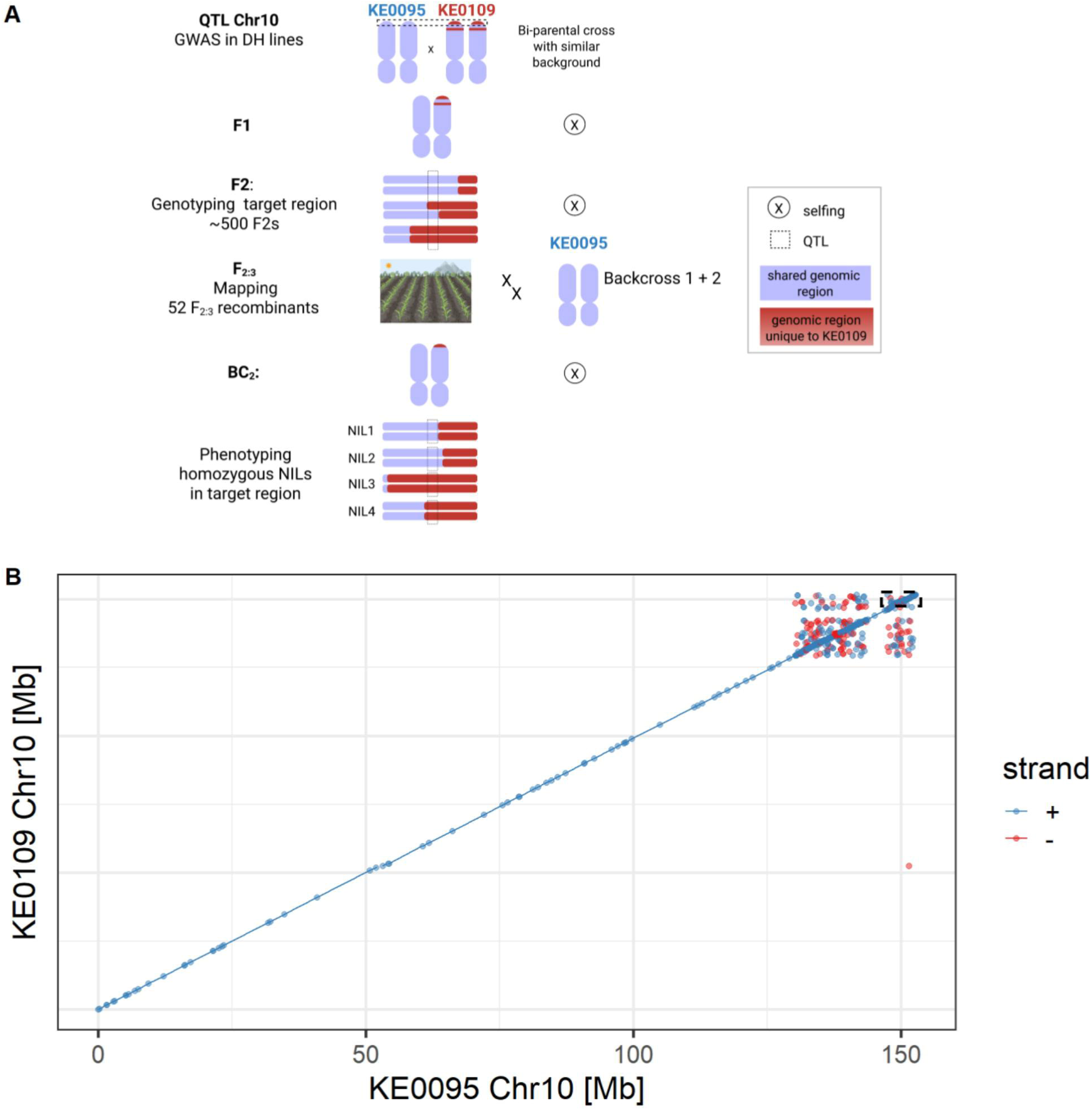
Material development for fine-mapping of a QTL for F_v_/F_m_ on chromosome 10. **A**) Schematic representation of the development of near isogenic lines segregating for a genomic region on chromosome 10 associated with F_v_/F_m_. Genomic fragments unique to the parental line KE0109 are highlighted in red. The QTL region is indicated by a dashed box. **B**) Pairwise sequence alignment of KE0095 and KE0109, the parents of the bi-parental population. The dashed box at the distal end of chromosome 10 marks the target region, which harbors the QTL for F_v_/F_m_. Alignments >1 kb with sequence identity >95% are shown: blue represents alignments on the forward strand, red indicates alignments on the reverse strand. Figure 2A was created with BioRender.com.

**Figure 3.**
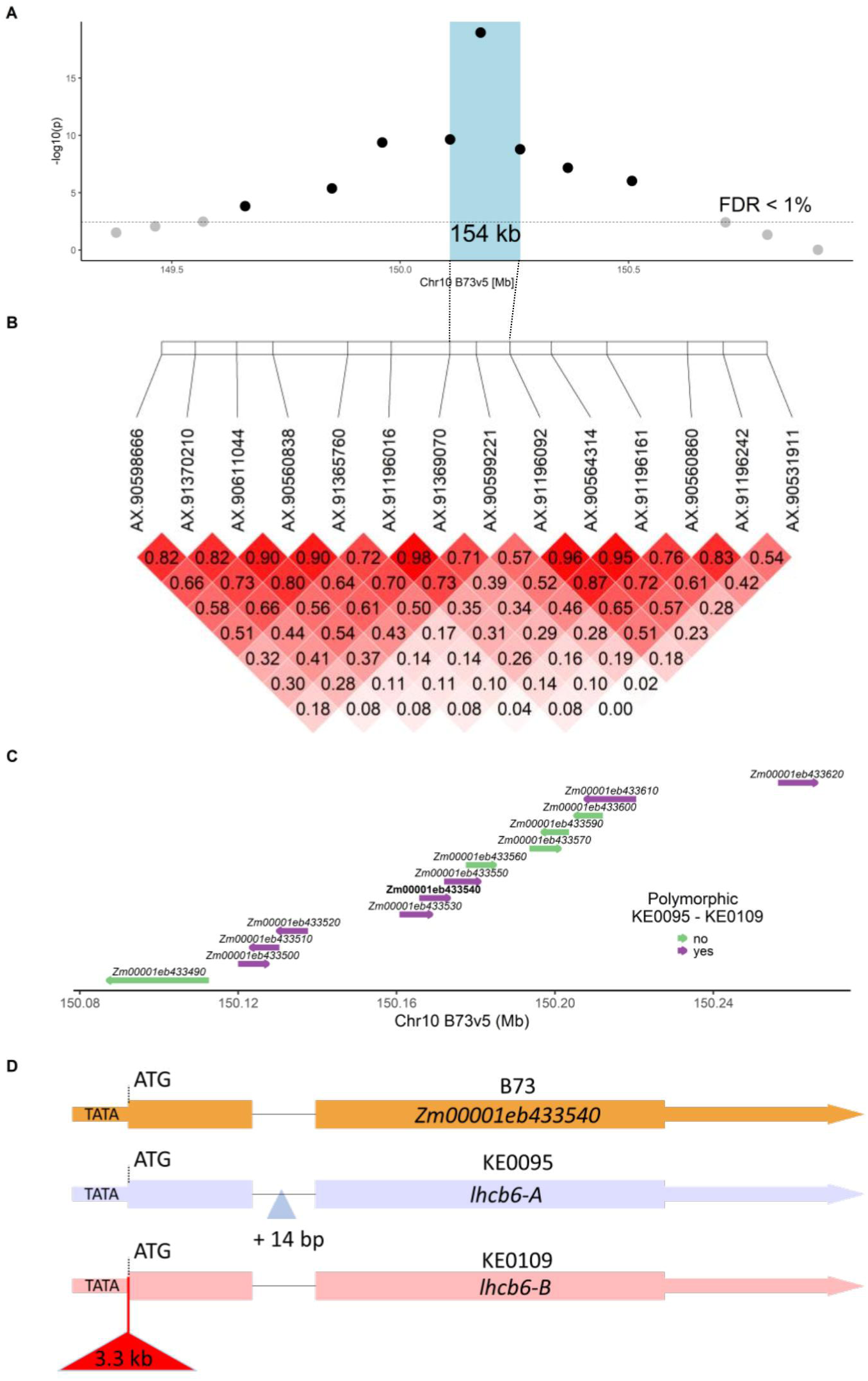
Characterization of a QTL region associated with F_v_/F_m_ in F_2:3_ recombinant lines. **A**) Association of 14 SNP markers with F_v_/F_m_ in F_2:3_ recombinant lines (N = 52) based on adjusted entry means across three field locations. The horizontal line indicates the false discovery rate (FDR) of 1%. The region between the flanking markers of the marker with highest significance is indicated by a blue-shaded box. **B**) Linkage disequilibrium (r^2^) between 14 SNPs in the F_2:3_ recombinant lines. **C**) Gene models located between the flanking markers of the marker with highest significance. The direction of the arrows indicates the read strand on which the respective gene model is located. Genes with polymorphisms between KE0095 and KE0109 are shown in purple, and monomorphic genes between KE0095 and KE0109 are shown in green. **D**) Gene models of the gene *light harvesting chlorophyll a/b binding protein* 6 (*lhcb6*) in B73 (*Zm00001eb433540*), KE0095 and KE0109. KE0109 carries a 3.3 kb genomic insertion immediately upstream of the start codon (red triangle).

### Molecular characterization of candidate genes

In the 154 kb region of interest, 13 genes are located in the B73v5 reference genome (Figure 3C, Table S2). Protein sequences of these genes were annotated and classified using MapMan4 (Schwacke et al. 2019), and proteins belonging to category “Photosynthesis” were selected. One gene met the selection criterium: *Zm00001eb433540 (light harvesting chlorophyll a/b binding protein 6; lhcb6*). Since annotation alone reveals only genes with known functions, candidate gene prioritization was also conducted by comparing genomic sequences of KE0095 and KE0109.

Of the 13 genes in the target region, eight genes showed polymorphisms between KE0095 and KE0109, with six being non-synonymous leading to amino acid exchanges (Figure 3C, Table S3). Amino acid exchanges were non-conservative in two of these genes: Zm00001eb433520 and Zm00001eb433550. The former encodes an uncharacterized protein and differs by multiple indels and four non-conservative amino acid exchanges between KE0095 and KE0109. The latter encodes PROTEIN S-ACYLTRANSFERASE38 and exhibits one non-conservative amino acid exchange (Table S3). In addition, a 3.3 kb insertion directly upstream of the *lhcb6* translational start codon was detected in KE0109 which is not present in KE0095 and maize reference line B73 (Figure 3D, File S1). The corresponding 5’ sequence of *lhcb6* in B73 and KE0095 contains a sequence resembling a TATA box (TATTATTATTAAT; chr10:150,168,719-150,168,732; B73v5, Figure 3D, Figure S3) and is highly conserved across 33 maize genomes (Figure S4).

As a next step we investigated differences in transcript and protein abundance in leaves. In leaf proteomes of KE0095 and KE0109, 6,672 proteins were quantified. Of the 13 candidate genes two were represented (*Zm00001eb433540* [*lhcb6*], *Zm00001eb433610* [*chlh1*], Table S2). For *Zm00001eb433610* no difference in protein accumulation was observed between KE0095 and KE0109 (Table S2), while protein accumulation of LHCB6 was reduced ∼200 fold in KE0109 compared to KE0095 (Figure 4B). The absence of the other 11 candidates in leaf proteomes of KE0095 and KE0109 cannot rule them out as candidate genes for the QTL but makes them less likely. For the two genes with non-synonymous amino acid exchanges (*Zm00001eb433520*, Zm00001eb433550) and *lhcb6*, transcript abundance was measured by RT-qPCR. No differences in transcript levels of *Zm00001eb433520* and *Zm00001eb433550* were observed between KE0095 and KE0109 (Figure S5), while a 1000- fold reduction of transcript levels of *lhcb6* was detected in KE0109 compared to KE0095 (Figure 4A).

**Figure 4.**
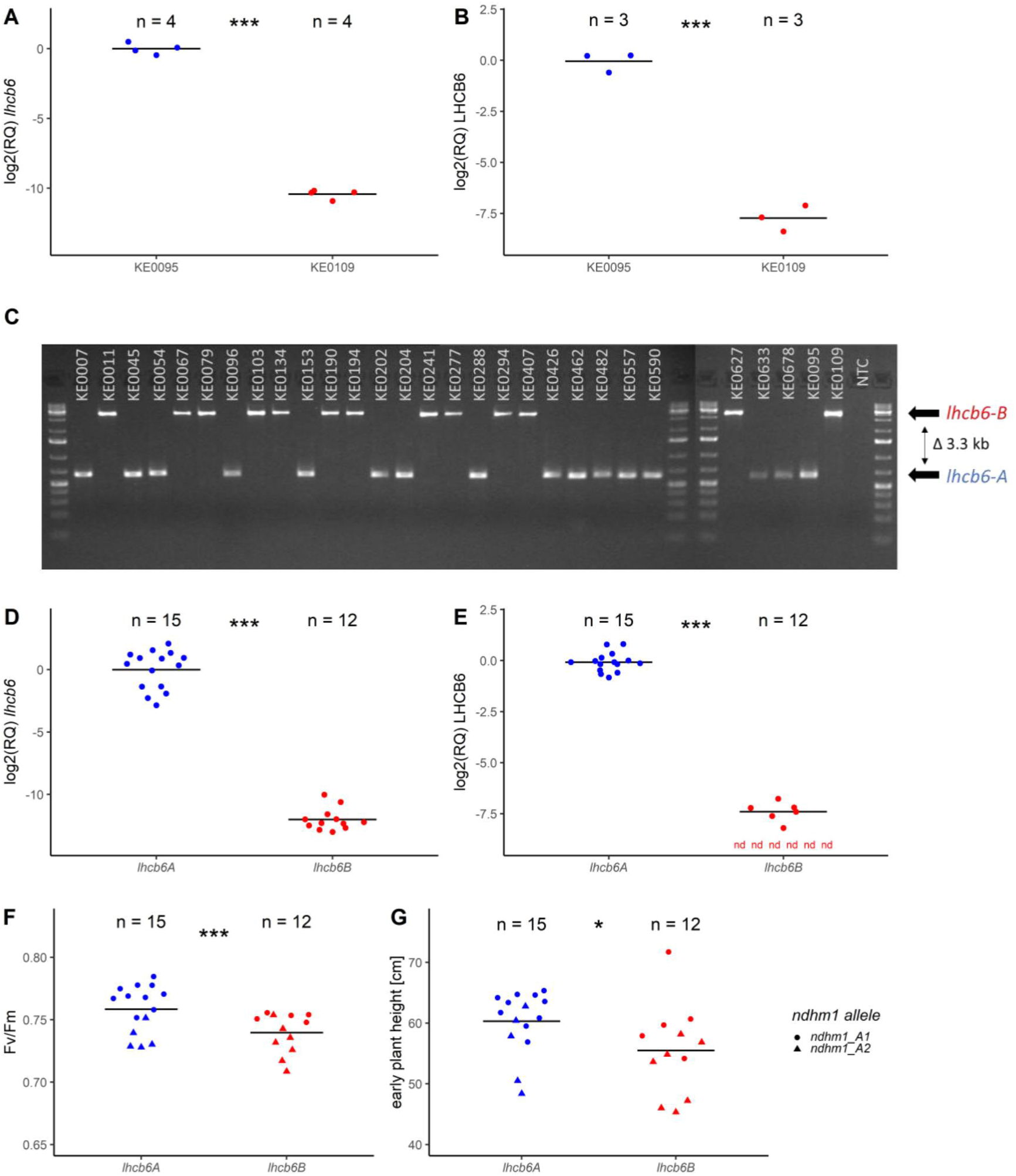
Molecular characterization of the candidate gene *lhcb6*. **A, B**) Log2-transformed relative transcript (**A**) and protein (**B**) levels of *lhcb6* in KE0095 and KE0109, normalized to KE0095. **C**) Genotyping of a 3.3 kb insertion in the genomic region upstream of the *lhcb6* start codon, assessed in a set of 27 Kemater lines and the parental lines KE0095 and KE0109. The lower band in the gel corresponds to the *lhcb6-A* allele and the higher band to the *lhcb6-B* allele, which contains the 3.3 kb genomic insertion. NTC: no template control **D-G**) Differences in transcript (**D**) and protein (**E**) abundance, F_v_/F_m_ **(F)** and early plant height **(G)** between 27 Kemater lines with contrasting *lhcb6* alleles. *lhcb6* (*lhcb6-A*, n = 15; *lhcb6-B*, n = 12) and *ndhm1* (*ndhm1-A1*, n = 15; *ndhm1-A2*, n = 12) segregate in the 27 Kemater lines. Genotypic data for *ndhm1* were derived from Urzinger et al. (2025). Significance was assessed by a two-way analysis of variance (ANOVA) including *lhcb6*, *ndhm1* and their interaction as factors. In **F** and **G** the *lhcb6* allele is indicated by color and *ndhm1* allele by shape of data points. Horizontal bars indicate group means and dots individual plant observations. Statistical differences were assessed by analyses of variance (ANOVA). ***P < 0.001; *P < 0.05. In **E** nd stands for not detected.

Assuming that sequence variation at *lhcb6* determines differences in F_v_/F_m_, we further characterized variation at *lhcb6*. The 3.3 kb insertion is highly similar to a putative hAT transposon that is located approximately 600 kb downstream of *lhcb6* in B73 according to MaizeGDB (File S2, DTA_ZM00039). The insertion contains two long open reading frames (313 and 492 amino acids), encoding domains typically found in transposases - a TTF zinc finger domain, a Ribonuclease H-like superfamily domain, and a hAT C-terminal dimerization domain (InterPro accessions: IPR006580, IPR008906 and IPR012337). However, no terminal inverted repeats at the insertion site, which are characteristic for hAT transposons were detected in KE0109. Further, KE0095 contains a 14 bp insertion within the only intron of *lhcb6* compared to both B73 and KE0109 (Figure 3D, File S1). Despite these differences in their DNA sequences, the amino acid sequences of LHCB6 in KE0095, KE0109, and B73 are identical. Hereafter, the KE0095 allele is referred to as *lhcb6-A* and the KE0109 allele with the 3.3 kb insertion is referred to as *lhcb6-B*.

Recently, *lhcb6* was proposed as a candidate gene for a QTL for F_v_/F_m_ in a maize MAGIC population, with maize line F7 carrying a deficient allele at this locus (Ferguson et al. 2025). We compared the genomic sequences of F7, KE0095 and KE0109 and found that F7 carried an identical insertion in its putative promotor sequence of *lhcb6* as KE0109, corroborating an effect of the insertion on *lhcb6* transcript levels in a different genomic background.

To investigate if the variation in *lhcb6* transcript and protein levels is specific to KE0095 and KE0109, 27 Kemater lines were selected from the population of DH lines used in GWAS analyses for validation experiments. The DH lines were chosen to represent the two alleles of the lead SNP of the F_v_/F_m_ QTL on chromosome 10 in equal proportion. The presence or absence of the *lhcb6-B* allele was determined by PCR analysis, with primers spanning the putative transposon insertion. The *lhcb6-B* allele was detected in 12 of the 27 Kemater lines and was in complete LD with the lead SNP from the GWAS in this set of Kemater lines (Figure 4C, Table S4). Corroborating results obtained with the two Kemater lines KE0095 and KE0109, RT-qPCR analysis revealed that *lhcb6* transcript levels were strongly decreased in the 12 Kemater lines carrying *lhcb6-B* compared to the 15 lines carrying *lhcb6-A* (Figure 4D).

Correspondingly, of the 12 lines with *lhcb6-B*, LHCB6 protein could either not be detected (N = 6) or was strongly reduced (N = 6) compared to the 15 lines with *lhcb6-A* (Figure 4E).

The 27 Kemater lines under study also harbor different alleles at *ndhm1*, a gene which was shown to affect F_v_/F_m_ and early plant height in maize (Urzinger et al., 2025). As expected from GWAS findings and results from F_2:3_ recombinant lines, mean F_v_/F_m_ was significantly (p < 0.01) decreased in DH lines with reduced LHCB6 protein levels in growth stage V6 (Figure 4F, Table S5). Further, early plant height in growth stage V6 was also significantly (p = 0.02) reduced in DH lines carrying the *lhcb6-B* allele (Figure 4G, Table S5). These results suggest that the transposon insertion in *lhcb6-B* diminishes *lhcb6* transcript abundance, leading to lower LHCB6 protein levels and thereby lower F_v_/F_m_.

To determine whether *lhcb6-B* is specific to the Flint maize heterotic group, haplotype diversity at this locus was investigated in 136 diverse inbred maize lines analyzed by Unterseer et al. (2016) and 501 DH lines derived from the Kemater landrace (Mayer et al., 2022). Haplotypes were defined by concatenating the 15 SNPs closest to *lhcb6* (8 upstream; 7 downstream). We identified 19 haplotypes with absolute counts ranging from 3 to 237 in the entire dataset. The most common haplotype (Hap1) had frequencies of 45% and 21% in the Kemater landrace and Flint breeding lines, respectively, but was absent in elite Dent lines (Figure S6). Based on the results of PCR genotyping, Hap1 was in complete LD with the insertion in *lhcb6-B* in the subset of 27 Kemater lines (Table S4).

### Functional characterization of the *lhcb6-B* allele

To evaluate the effect of the *lhcb6-A* and *lhcb6-B* alleles on phenotypic traits other than F_v_/F_m_, we developed four near isogenic lines with contrasting *lhcb6* alleles from the F_2:3:_ recombinant lines. NIL 1 and NIL 2 differ from NIL 3 and NIL 4 in a 154 kb genomic fragment harboring *lhcb6* and 12 other genes defined by fine-mapping in F_2:3_ recombinant lines (Figure 3). NIL 1 and NIL 2 carry *lhcb6-A*, while NIL 3 and NIL 4 carry *lhcb6*-*B* (Figure S7, Table S6). In addition to F_v_/F_m_, we measured the effective PSII antenna size, NPQ dynamics, and fresh and dry biomass of the NILs. As expected, F_v_/F_m_ was lower in NILs carrying *lhcb6-B* compared to NILs carrying *lhcb6-A* (Figure 5A), which was the result of a combination of higher minimal fluorescence (F_0_; Figure S8A) and lower maximal fluorescence (F_m_; Figure S8B). Effective antenna size of PSII was determined from the half time needed to reach the J level of the initial O-J rise in an O-J-I-P chlorophyll induction curve (Strasser et al., 2004). The half-time was significantly lower in KE0109 and NILs carrying the *lhcb6-B* allele indicating larger effective antenna size (Figure 5B). Maximum NPQ in NILs with *lhcb6-B* and KE0109 was approximately half of that observed in NILs with *lhcb6-A* and KE0095 (Figure 5C). Conversely, NPQ values during relaxation in the dark were increased in NILs with *lhcb6-B* and KE0109 (Figure 5C), suggesting impaired dynamic regulation of NPQ. Fresh and dry biomass were reduced in NILs with *lhcb6-B* compared to NILs with *lhcb6-A*, but this reduction was not observed in KE0109 (Figure 5D, E).

**Figure 5.**
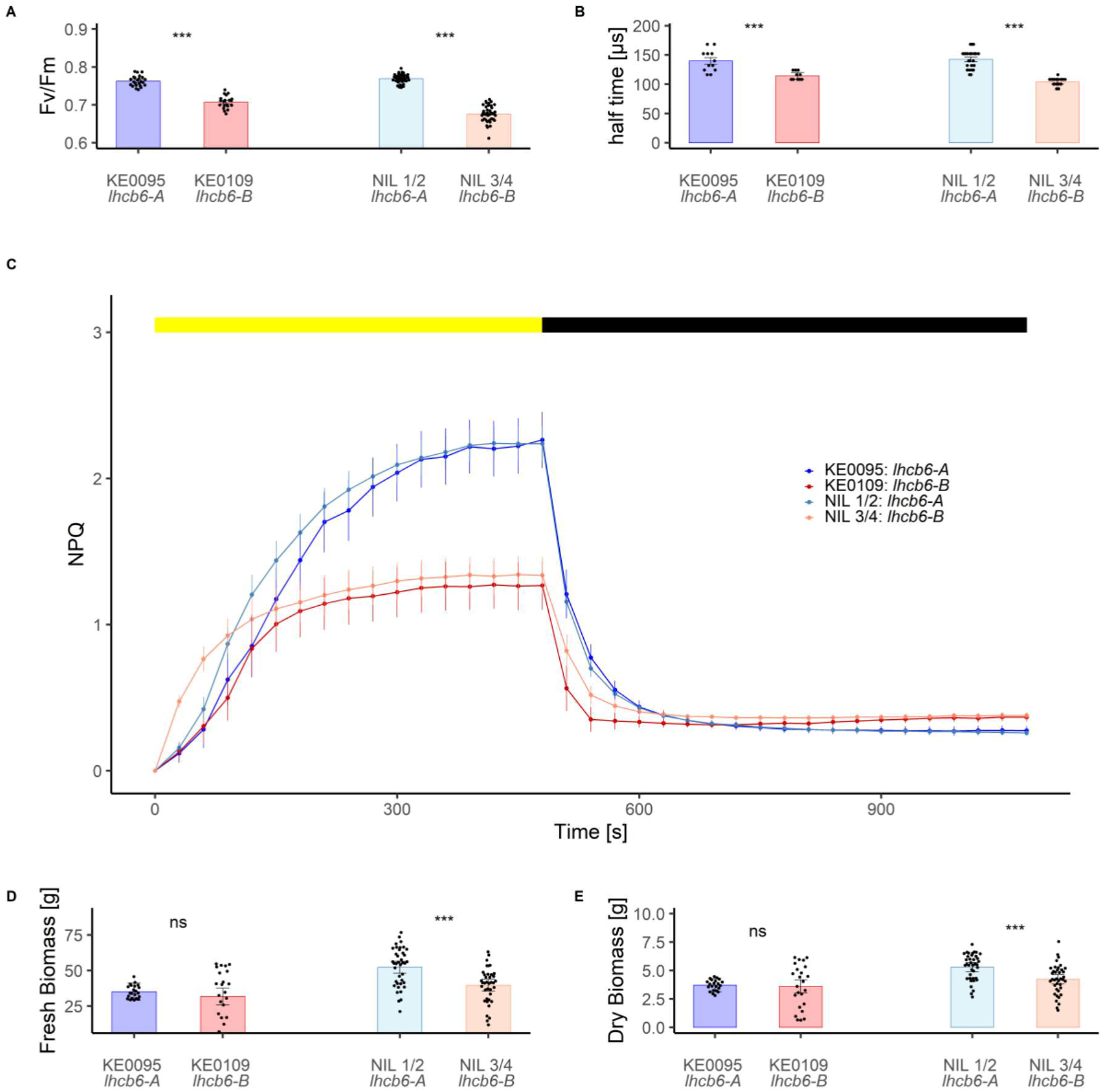
Physiological and agronomic evaluation of near isogenic lines (NILs) with contrasting *lhcb6* alleles and of parental lines KE0095 and KE0109. Plants were grown in a growth chamber under fluctuating light intensity until growth stage V6. **A**) Maximum quantum yield of photosystem II (F_v_/F_m_). **B**) Functional antenna size estimation based on the half-time to the J level (T₁/₂) during an OJIP fluorescence induction curve. **C**) Non-photochemical quenching (NPQ) dynamics during an eight-minute light phase (yellow bar) followed by a ten-minute dark phase (black bar). Dots represent adjusted means, bars represent standard errors (SE) **D)** Fresh biomass. **E**) Dry biomass. Measurements were taken at growth stages V5–V6. Significant differences based on Student’s t-tests are indicated by stars. ***P < 0.001; ns: not significant. Bars represent adjusted means ± SE and dots represent individual plant observations (**A**, **B**, **D**, **E**).

We further explored the consequences of reduced *lchb6* expression on LHCII antenna components in the subset of 27 Kemater lines described earlier. Based on sequence similarity to experimentally annotated *Arabidopis thaliana* LHCII components, 13 putative LHCII subunits were detected in the maize reference line B73 (Table S7). One of these is an uncharacterized isoform of LHCB6 (Zm00001eb066480, LHCB6-2). LHCB6-2 has an amino acid sequence identity (235/253 amino acids) with LHCB6-1, including a four amino acid deletion at the N- terminus in LHCB6-2 compared to LHCB6-1. Of these 13 proteins, 12 were detected and quantified in the leaf proteomes of the 27 Kemater lines, including the three major LHCII antenna proteins (LHCB1, LHCB2 and LHCB3) and the three minor LHCII antenna proteins (LHCB4, LHCB5 and LHCB6). In addition to the expected reduction in LHCB6, LHCB3 (*Zm00001eb324240*) protein levels were decreased by ∼50% in *lhcb6-B* lines. The abundance of the other 10 quantified LHCII proteins, including LHCB6-2, remained unchanged. We also analyzed differential accumulation among all 5,531 quantified proteins. Only six proteins exhibited differential accumulation based on *lhcb6* allele status (Table S8, Figure 6). The *lhcb6- B* allele was correlated with a reduced abundance of four proteins and an increased abundance of two proteins (Figure 6, Table S8). Notably, LHCB6 showed the most significant reduction, followed by LHCB3 (Figure 6A).

**Figure 6.**
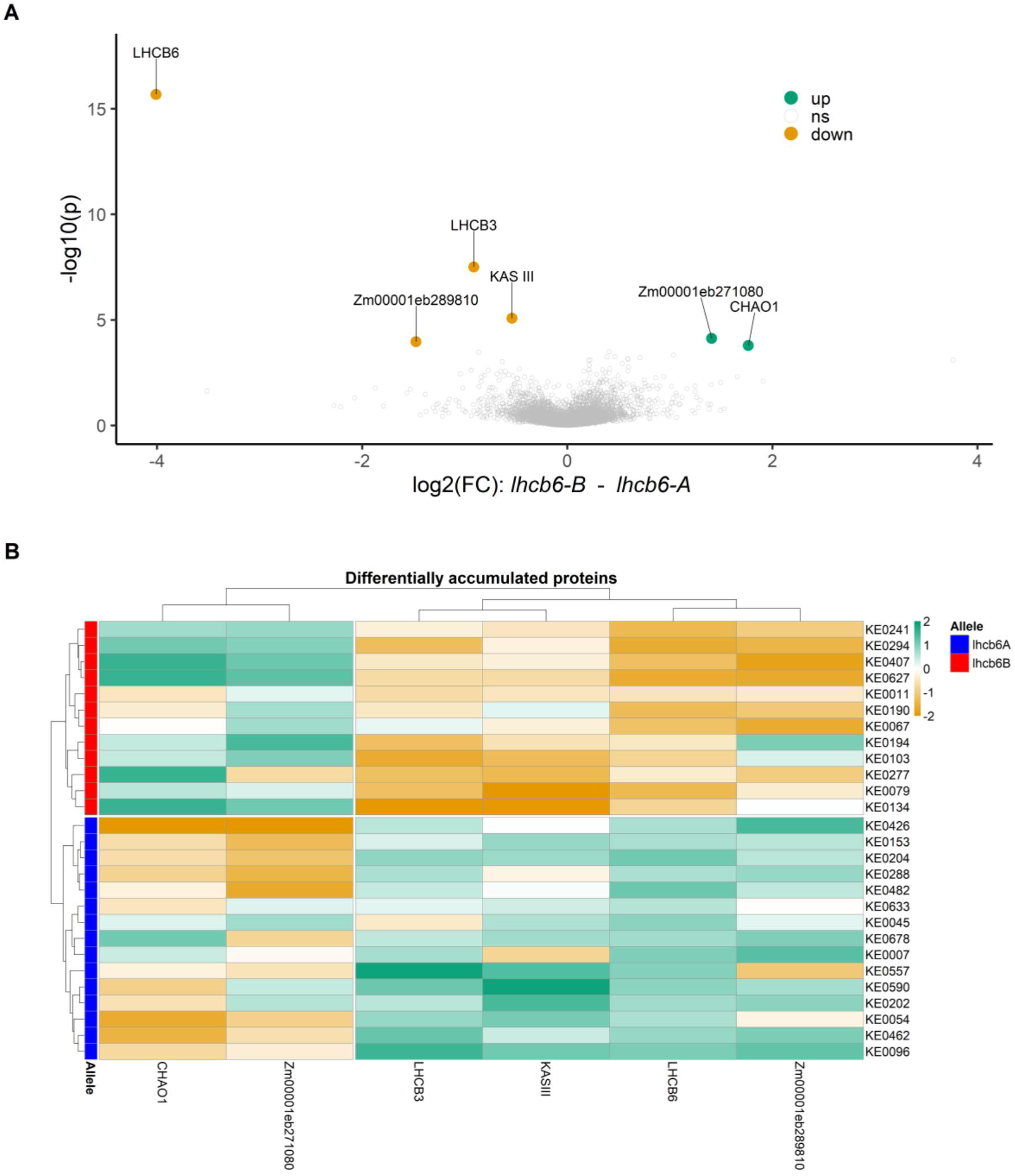
Leaf proteome analysis of 27 Kemater lines segregating for *lhcb6-A* and *lhcb6-B*. **A**) Differentially accumulated proteins in 27 Kemater lines grouped by their *lhcb6* allele. Proteins were considered differentially accumulated at a false discovery rate (FDR) < 20%. Proteins with lower abundance in lines carrying the *lhcb6-B* allele are shown in orange, while those with higher abundance are shown in green. **B**) Relative quantities of six significantly differentially accumulated proteins from (**A**) expressed as z-scores for each of the 27 Kemater lines. Lines carrying the *lhcb6-A* allele indicated by the blue and those with the *lhcb6-B* allele with the red bar.

Of the two other proteins with decreased accumulation in Kemater lines with *lhcb6-B* (Zm00001eb068760, Zm00001eb289810), only one has a clearly defined function: an isoform of 3-oxoacyl-[acyl-carrier-protein]-synthase III (KAS III; Zm00001eb068760). KAS III catalyzes the first step in the fatty acid synthesis pathway II, which is essential for the synthesis of membrane phospholipid acyl chains (White et al., 2005).

Among proteins with increased abundance, CHLOROPHYLLIDE A OXYGENASE1 (CHAO1; Zm00001eb039350) showed the strongest upregulation — a threefold increase in lhcb6-B lines (Figure 6B). Another protein with increased accumulation is of unknown function (Zm00001eb271080).

## Discussion

### Unraveling the genetic basis of photosynthetic efficiency in maize

A profound understanding of the genetic basis of photosynthetic efficiency can leverage the improvement of agricultural productivity (Kromdijk et al., 2016, Theeuwen et al., 2022, Leister, 2023). Previous research has established a genetic correlation between the maximum quantum yield of PSII (F_v_/F_m_) and agronomically relevant traits like early development and chilling stress tolerance in maize, and links it to other photosynthetic traits, including PSII operating efficiency (ΦPSII), non-photochemical quenching (NPQ), and chlorophyll content (Hölker et al., 2019, Lainé et al., 2023, Urzinger et al., 2025, Ferguson et al., 2025). QTL with large effects on F_v_/F_m_ have been mapped in maize and several candidate genes have been proposed, but their underlying genetic mechanisms have remained largely unknown (Fracheboud et al., 2002, Fracheboud et al., 2004, Hund et al., 2004, Jompuk et al., 2005, Ferguson et al., 2025).

In this study, we identified five QTL for F_v_/F_m_ across two early growth stages segregating in a maize landrace. The causal gene underlying the QTL on chromosome 2 (*ndhm1*) with pleiotropic effects on early plant growth and F_v_/F_m_ has been characterized by our group recently and encodes a subunit of the NADH-dehydrogenase like (NDH) complex (Urzinger et al., 2025). Here, we could show that *lhcb6* (also known as CP24), a minor antenna component of the light-harvesting complex II (LHCII), is the causal factor underpinning the major QTL for F_v_/F_m_ on chromosome 10. Both *lhcb6* and *ndhm1* play key roles in dynamic regulation of photosynthesis, by balancing photoprotection and efficiency via NPQ and cyclic electron transport, respectively (Murchie and Ruban, 2020, Ishikawa et al., 2016). We therefore conclude that F_v_/F_m_ can serve as a proxy trait for identifying meaningful genetic variation that contributes to photosynthetic regulation in changing environmental conditions.

### Functional characterization of lhcb6 as a major determinant of Fv/Fm in maize

In a recent publication, Ferguson et al. (2025) proposed *lhcb6* as candidate gene for a QTL affecting F_v_/F_m_, ΦPSII and NPQ in a maize multiparent advanced generation inter-cross (MAGIC) population. They showed that a specific haplotype found in the Flint line F7 was associated with reduced transcript and protein levels of *lhcb6* compared to the maize reference line B73. Our current study detected a QTL in the same genomic region in a flint maize landrace. Ferguson et al. (2025) based their evidence for *lhcb6* being the causal gene on functional annotation, physiological similarity to Arabidopsis (*Arabidopsis thaliana*) mutants, and expression differences between B73 and F7, leading to impaired assembly of higher order PSII-LHCII supercomplexes. By mapping short-read sequencing data from F7 against the B73 reference, they identified three SNPs and one indel specific to the F7 haplotype altering computationally predicted transcription factor binding sites (TFBS). Here, we used a forward genetic approach with near-isogenic lines (NILs) derived from a maize landrace in combination with long-read genome assemblies of their two parental lines KE0095 and KE0109 and we could pinpoint the causal polymorphism underlying this QTL to a 3.3 kb insertion of a hAT transposon directly upstream of the start codon of *lhcb6* in the *lhcb6-B* allele. The functional importance of the affected region as promotor of *lhcb6* is supported by its conservation across a diverse panel of maize lines, its overlap with a region of accessible chromatin, indicating its accessibility to transcription factors (Ricci et al., 2019), computationally predicted TFBS (Ferguson et al., 2025), and a putative TATA-box. The insertion of a hAT transposon in *lhcb6- B* shifts the putative promotor region of *lhcb6* 3.3 kb upstream, extending the typical distance of *cis-*regulatory elements to the transcription start site in maize (Savadel et al., 2021). We also identified the same mutation in the Flint line F7, linking our findings with those of Ferguson et al. (2025). Thus, our results strongly support the hypothesis that *cis*-genic allelic variation at *lhcb6* is causative for differential gene expression, leading to significantly reduced RNA and protein levels, and further enabled us to identify the specific underlying structural variation.

In most vascular plants, the LHCII antenna consists of major antennae (homo- or heterotrimers of LHCB1, LHCB2, and LHCB3) and minor antennae (monomers of LHCB4, LHCB5, and LHCB6) (Ballottari et al., 2012, Iwai et al., 2024). These antennae assemble into PSII supercomplexes (e.g., C_2_S_2_M_1_ C_2_S_2_M_2_, C_2_S_2_), which are composed of two core (C) subunits of PSII, two strongly (S) bound trimers, and zero, one, or two moderately (M) bound trimers. In these supercomplexes, the M trimer is coordinated by LHCB3 and LHCB6 (Su et al., 2017). Our proteome analyses show both significantly reduced LHCB6 and LHCB3 levels in DH lines carrying the *lhcb6-B* allele reflecting the direct interaction between the two proteins. A lack of LHCB6 protein impairs LHCII assembly, favoring the formation of C_2_S_2_ complexes (Kovacs et al., 2006). This can explain the raised F_0_ levels we observed in NILs due to reduced connectivity between antennae and the PSII core (Albanese et al., 2016). Interestingly, although *lhcb6* mutants severely impair growth in Arabidopsis, the *lhcb6-B* allele in maize causes only mild growth reductions, suggesting compensatory mechanisms (Ganeteg et al., 2004, Kovacs et al., 2006, De Bianchi et al., 2008). In addition, the unexpected high frequency of the *lhcb6-B*-associated haplotype in Flint breeding lines certainly warrants further research on the role of LHCB3 and LHCB6 in maize photosynthesis. Though LHCB6 and LHCB3 were once thought essential in land plants, some gymnosperms lack them entirely (Kouřil et al., 2016). Thus, whether the Flint specificity of *lhcb6-B* reflects selection for temperate climates, as proposed by Ferguson et al. (2025), or random drift is unclear and requires further investigation

### Effects of *lhcb6-B* on the maize photosynthetic machinery

LHCII plays a dual role in light harvesting and photoprotection. It captures and transfers light energy to PSII and is involved in photoprotection through NPQ, a mechanism that dissipates excess light energy as heat (Murchie and Ruban, 2020). Genetic variation for NPQ has been reported for maize and sorghum (*sorghum bicolor*) populations (Sahay et al., 2023, Sahay et al., 2024). In maize, LHCII regulation responds to light and temperature (Caffarri et al., 2005), with implications for state transitions and energy balance between the two photosystems (Kargul and Barber, 2008, Pan et al., 2018). Thus, targeting LHCII dynamics is a potential strategy to enhance stress resilience (Kromdijk and Walter, 2023) and maize landraces can enhance our understanding of these complex physiological processes (Liang et al., 2021, Theeuwen et al., 2022).

Employing genomic and proteomic data from our landrace population we investigated potential compensatory mechanisms related to *lhcb6-B* in maize. Our analysis revealed a previously uncharacterized isoform of LHCB6 in maize, which we designated as LHCB6-2 (*Zm00001eb066480*). While isoforms for LHCB1, LHCB2, and LHCB4 have been reported in Arabidopsis (Cheng et al., 2017), this is, to our knowledge, the first description of a LHCB6 isoform. The two maize LHCB6 isoforms are highly similar (File S3), with most differences occurring at the N-terminus, a region that includes a potential phosphorylation site (Iwai et al., 2024). Although phosphorylation is known to regulate LHCB proteins in species such as spinach (*Spinacia oleracea*), Arabidopsis, and *Chlamydomonas reinhardtii*, it has not been documented for LHCB6 (Pan et al., 2021, Dunahay et al., 1987). As LHCB6-2 cannot compensate for the deficiency of LHCB6-1 (*Zm00001eb433540*) in the Kemater DH lines we propose that LHCB6-1 and LHCB6-2 may have distinct roles. While LHCB6-1 is important for photoprotection balanced by NPQ, LHCB6-2 may primarily facilitate efficient light capture for photosynthesis, partially compensating for negative effects on biomass accumulation. However, this hypothesis requires further analyses which are beyond the scope of this study.

Comparing the leaf proteome between contrasting Kemater lines carrying either the *lhcb6-A* or the *lhcb6-B* alleles, we identified several differentially accumulated proteins shedding light on the dynamic regulation of the LHCII antenna. For instance, chlorophyllide a oxygenase (CHAO1; also known as CAO), which converts chlorophyll a to chlorophyll b and is essential for LHCII antenna assembly (Tanaka and Tanaka, 2011), shows elevated levels in lines carrying *lhcb6-B*. Increased CHAO1 may serve as a compensatory feedback mechanism in response to LHCII antenna deficiency. A connection between chlorophyll *b* levels and LHCII efficiency has been shown in other species. In Arabidopsis and barley (*Hordeum vulgare*), deficiencies in chlorophyll *b* have been shown to lead to reduced accumulation of LHCII components (Havaux et al., 2007, Bossmann et al., 1997), and overexpression of CHAO1 in Arabidopsis enlarged the LHCII antenna (Tanaka et al., 2001). Further, an uncharacterized protein potentially involved in lipid transfer (*Zm00001eb271080*) has been linked to variation in maize tocopherol and tocotrienol content (Diepenbrock et al., 2017). Given the protective role of tocopherols against photoinhibition and lipid photooxidation (Havaux et al., 2005), its increased protein levels might indicate increased oxidative stress associated with impaired NPQ, similar to the increased reactive oxygen species (ROS) observed in Arabidopsis *lhcb6* mutants (Chen et al., 2018). Reduced protein levels of KAS III, an enzyme involved in lipid biosynthesis, may reflect disturbances in the assembly of the PSII–LHCII supercomplex, where lipids play a stabilizing role (Sheng et al., 2018). The exact mechanisms underlying the proteomic plasticity associated with the *lhcb6-B* protein will be further investigated.

## Conclusion

We identified and characterized a major QTL for F_v_/F_m_ in a maize landrace, attributing it to a hAT transposon insertion in the *lhcb6* promoter. This mutation drastically reduces LHCB6 expression, disrupts LHCII antenna assembly, and affects the photoprotective mechanism NPQ. Proteomic analyses further indicate that compensatory responses aimed at maintaining photosynthetic efficiency occur in the maize leaf. Our findings highlight the value of natural genetic variation for unraveling the complex regulation of photosynthesis and for enhancing crop performance under stress conditions.

The work presented here also has significant implications for breeding. The development of a diagnostic marker capturing the allelic state of *lhcb6* - including the hAT transposon insertion in the promotor - and using this marker alone or in combination with two or more flanking markers to define a diagnostic haplotype will facilitate the evaluation of the gene’s effect in segregating populations and enable any type of marker-based selection including whole-genome selection. In addition, the screening of additional natural genetic variation and the generation of new variation at *lchb6* via chemical induced mutations, genome editing, or classical transgenic approaches may allow engineering of the LHCII antenna and modification of NPQ in breeding material. Beneficial alleles detected or created in this manner can be introgressed into elite material through marker assisted backcrossing.

## Material and Methods

### Plant Material

The landrace “Kemater Landmais Gelb” (Kemater) was chosen as the base population for this study due to its phenotypic variation for early development, as well as its fast decay of linkage disequilibrium (LD) and absence of pronounced population structure. Kemater was selected from a set of 35 European maize landraces covering a broad geographical region described by Mayer et al. (2017). Kemater represents 77% of the molecular variance of the entire collection (Hölker et al., 2019) based on the 600k Axiom™ Maize Genotyping Array (600k Array, Unterseer et al., 2014). Directly from the landrace, 516 doubled-haploid (DH) lines were produced and multiplied using the in vivo haploid induction method (Röber et al., 2005), and phenotyped in 2017 and 2018. A curated dataset comprising 501 Kemater DH lines genotyped with 501,124 markers from the 600k Array was obtained from Mayer et al. (2022).

A subset of 27 DH lines was selected from the full set of Kemater lines for in-depth analysis of a QTL for F_v_/F_m_ on chromosome 10 (Chr10: 148,413,861-152,303,534; Table1, Figure 1). These lines were chosen to obtain a balanced frequency of two contrasting alleles at the QTL (validation set, Table S4).

### Development of recombinant and near isogenic lines

A schematic overview of the material development for fine mapping is presented in Figure 2A. Two Kemater DH lines, KE0095 and KE0109, were chosen for their contrasting alleles at the QTL for F_v_/F_m_ on chromosome 10, while being as similar as possible in their genomic background (Figure 2B, Figure S2) and crossed to produce a bi-parental population. KE0095 and KE0109 are polymorphic for 74,975 of 493,989 markers of the 600k Array (Unterseer et al., 2014). KASP markers polymorphic between the two parental lines and positioned in the QTL region on chromosome 10 were synthesized using probe sequences of the 600k Array. F_2_ lines were genotyped with six KASP markers and individual plants showing recombination between markers AX-90560736 (Chr10:143,112,717; Table S1) and AX-91196698 (Chr10:152,357,073; Table S1) were self-pollinated. Resulting F_2:3_ recombinant lines were genotyped with 14 KASP markers (Table S1) and phenotyped for F_v_/F_m_ in field experiments at three German locations in 2024. Selected F_2:3_ recombinants were backcrossed two times to KE0095 as the recurrent parent. BC_2_ was genotyped with a 15k SNP chip to select individual plants with minimum donor parent genome. These were selfed to obtain four near-isogenic lines (NILs) which were phenotyped in growth chamber experiments.

### Field experiments

The Kemater DH lines were evaluated in 2017 and 2018 as part of a broader study on landrace diversity (Hölker et al., 2019). Briefly, in two locations, Golada and Tomeza, 222 Kemater DH lines were phenotyped for F_v_/F_m_. The experimental design was a 10 × 10 lattice design with two replications at each site. As checks, fourteen Flint and one Dent inbred line as well as the original landrace population (each replicated four times) were used. Further information on field trial locations, check varieties, and methods for phenotypic data analysis is available in the publications of Hölker et al. (2019) and Mayer et al. (2022).

Phenotypic evaluation of the set of 27 Kemater DH lines and the population of recombinant F_2:3_ lines used for fine-mapping was conducted in Einbeck (EIN, Germany, 51°49’05.9"N 9°52’00.3"E), Roggenstein (ROG, Germany, 48°10’47.5"N 11°19’12.9"E), Bernburg (BBG, Germany, 51°49’28.6"N 11°42’26.3"E) and Freising (FRS, Germany, 48°24’13.2"N 11°43’28.5"E). The subset of 27 Kemater DH lines was grown in a randomized complete block design (RCBD) with two replications in locations ROG, FRS and Oberer Lindenhof (OLI, Germany, 48°28’26.3"N 9°18’17.9"E) in 2023 together with four inbred checks. The 52 F_2:3_ recombinant lines were evaluated in a RCBD with two replications in EIN, ROG and FRS in 2024 using the same inbred checks. Plots consisted of 20 plants grown in single rows of 3 m length with 0.75 m spacing between them (9 plants/m^2^). Field trials were subjected to standard agricultural practices. Phenotypic observations from plots containing at least five plants were filtered by a Grubbs’ outlier test (Grubbs, 1969).

### Growth conditions in growth chamber experiments

For phenotyping of the four NILs, alongside their recurrent parent (KE0095) and donor parent (KE0109) in growth chamber experiments, kernels were imbibed in water for 5 min, transferred to filter paper and pre-germinated for 72 h in the dark at 28 °C. After germination, plant material was sampled from individual seedlings for DNA extraction and the respective seedlings were transferred to small pots filled with CL ED73 soil (Einheitserdewerke Patzer, Germany). Plants were grown in growth chambers with 16/8 h day night [d/n], 25/20 °C d/n, 75% relative humidity [RH] until growth stage V4 to V6. To simulate fluctuations in light intensity occurring in the field, a light switch scheme was employed, alternating between 30 minutes high light (800 µmol m^-2^s^-1^ photosynthetically active radiation [PAR]) and 15 minutes low light (100 µmol m^-2^s^-1^ PAR). The four NILs and the two parental DH lines KE0095 and KE0109 were replicated 10 times in each of the two blocks in a RCBD. The developmental stages of maize were determined according to the leaf collar method (Ritchie et al., 2005). Phenotypic data from growth chamber experiments were filtered by a Grubbs’ outlier test (Grubbs, 1969).

### Measurement of photosynthetic traits

F_v_/F_m_ was measured on the last fully developed leaf in dark adapted plants before light was turned on in the growth chambers using a LI-600 porometer (LI-COR Inc., Lincoln, NE, USA). In field experiments, the last fully developed leaf was cut from the plant and dark adapted for at least 30 minutes before measuring. F_v_/F_m_ was assessed in growth stage V6 with a LI-6800 Portable Photosynthesis System (LI-6800; LI-COR Inc., Lincoln, NE, USA). The last fully developed leaf was clipped in the middle, sparing the midvein, flow rate set to ‘high’ (150 µmol*s^-1^), match frequency of 10, flash set to ‘dark adapted’ and ‘Rectangular’ with an intensity of 6000 µmol m^-2^s^-1^, a flash duration of 800 ms, and the fluorescence constants ‘Leaf absorptance’ and ‘Fraction Abs PSII’ to 0.8 and 0.5 respectively at a modulation rate of 5 Hz. Non-photochemical quenching (NPQ) was measured with a LI-6800. The maximum fluorescence F_m_ was determined on the last fully developed leaf, with the flash set to ‘Rectangular’ with an intensity of 6000 µmol m^-2^s^-1^, a flash duration of 800 ms, an output rate of 100 Hz, a margin of 5 points and dark mode rate of 50 Hz. Light adapted maximum fluorescence F_m_‘ was determined after setting the actinic light to 1500 µmol m^-2^s^-1^ by conducting 16 saturating flashes every 30 seconds from 30 up to 480 seconds after illumination and after subsequently switching off the actinic light by another set of 20 saturating flashes every 30 seconds from 30 to 600 seconds after dark. NPQ was calculated for each measurement timepoint separately as (F_m_ - F_m_‘) / F_m_‘. For estimation of PSII antenna size, fast chlorophyll a fluorescence induction kinetics (OJIP transients) were measured by a flash with intensity of 20,000 µmol m^-2^s^-1^ and a duration of 600 ms with a LI-6800. The Dark mode rate was set to 500 Hz, the Light mode rate to 1 kHz and the Flash mode rate to 250 kHz. The half rise time t1⁄2 from the minimal fluorescence F_0_ to the fluorescence at time point J (F_j_) was used as an estimate of the effective antenna size of PSII (Strasser et al., 2004).

### Plant height measurement in the field

Early plant height was measured by stretching all leaves of a plant to measure the maximum length between soil and the tip of the leaves in growth stages V4 and V6. The measurements of three individual plants per plot were averaged to obtain the plot level measurement.

### Identification of trait associations

We conducted a genome-wide association study (GWAS) for F_v_/F_m_ in growth stages V4 and V6 in the Kemater population with the software GEMMA (v 0.98.1) (Zhou and Stephens, 2012). Genotypic data for the 222 Kemater DH lines was obtained from Mayer et al. 2022 and filtered for polymorphic SNPs. Phenotypic data for F_v_/F_m_ estimates were obtained from Hölker et al. (2019). Briefly, F_v_/F_m_ was assessed in a set of Kemater DH lines (V4: n = 222; V6: n = 211) using a fluorometer (OS-30p, Opti-Sciences Inc., USA) at growth stages V4 in both 2017 and 2018, and V6 in 2017. The following statistical model was used:

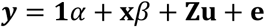

where **y** is the *n*-dimensional vector of adjusted entry means for F_v_/F_m_ averaged across four (V4) and two (V6) environments respectively, with *n* being the number of DH lines; α is the intercept; *β* is the fixed effect of the tested SNP; **x** is the vector of corresponding genotype scores coded as 0 or 2; **u** is the *n*-dimensional vector of random genotypic effects, with **u**∼*N*(**0**,**K***σ^2^*); and **e** is the *n*-dimensional vector of random residual effects, with **e**∼*N*(**0**,**I** *σ^2^*). **K** denotes the (*n* × *n*) genomic kinship matrix based on SNP markers, calculated with GEMMA (“-gk-2”). **I***_n_* denotes the (*n* × *n*) identity matrix. *σ^2^* and *σ^2^*refer to the genetic and residual variance pertaining to the model, respectively. The matrix **Z** (n × n) assigns adjusted means to the random genotypic effects. Significance of trait associations was assessed with a likelihood ratio test, as implemented in GEMMA, using a 15% false discovery rate (FDR; Benjamini and Hochberg, 1995). Pairwise r^2^ between SNPs was calculated as their squared pearson correlation coefficient (Hill and Robertson, 1968). SNPs with an r^2^ ≥ 0.6 and within a physical distance < 1 Mb were considered to belong to the same QTL. Physically overlapping QTL were merged and the resulting QTL was described by the start and end position of the first and last SNP from the merged QTL. The proportion of genetic variance explained by a single QTL was estimated by calculating the reduction in genetic variance between a full model including all QTL and a reduced model excluding the respective QTL, following Mayer et al. (2020).

### Transcript level measurements

RNA was extracted from leaves using a guanidine hydrochloride protocol (Logemann et al., 1987), followed by DNase digestion and first-strand cDNA synthesis (Maxima H Minus Kit, random hexamer primers, Thermo Scientific K1652). RT-qPCRs were performed in technical triplicates with both primers binding to the 3’ UTR of *lhcb6, Zm00001eb433520 and Zm00001eb433550* (Table S9). For normalization, RT-qPCR results of the house-keeping gene MEP served as reference (membrane protein PB1A10.07c, *Zm00001eb257640*).

### Leaf proteome analysis

For proteome measurements the parental lines KE0095 and KE0109 were grown in a growth chamber as described above. Leaf material was sampled in frozen nitrogen for three biological replicates per genotype in growth stage V4. From the subset of 27 DH lines grown in field experiments, a pooled sample of six plants from two replicates of one genotype was taken from the last fully developed leaf in growth stage V6 and frozen in liquid nitrogen. Total proteome was extracted and measured following an established protocol (Brajkovic et al., 2023). For quantification of peptides an additional labeling step was added after protein digestion. Samples of KE0095 and KE0109 were labeled using 16-plex tandem mass tag (TMT) reagent, while samples from the subset of 27 DH lines were labeled using 11-plex TMT reagent and processed in three batches. A detailed protocol for sample preparation and mass spectrometry was described by Urzinger et al. (2025). For normalization of batch effects in the subset of 27 Kemater lines, a reference was prepared by mixing equal peptide amounts of all samples which was measured twice in each of the three TMT batches. Proteins were identified using the proteome of B73v5 as reference (Hufford et al., 2021) and quantified by MaxQuant (Tyanova et al., 2016). To avoid imputation of missing values in quantitative analyses, only proteins which were identified in all samples of an experiment were considered. Log2 transformed raw intensities of identified proteins were median normalized to remove differences in summed intensities per sample. For the subset of 27 Kemater lines the two reference channels per TMT batch were used to calculate a correction factor for each identified protein per TMT batch.

### Detection of the *lhcb6-B* allele

To determine presence or absence of a 3.3 kb insertion in the genomic sequence upstream of the *lhcb6* start codon (Figure 3D) a PCR assay was developed. Primer sequences are listed in Table S9. PCR reactions were performed using Q5 High-Fidelity DNA Polymerase (New England Biolabs Inc., Ipswich, MA, USA) according to the manufacturer’s specifications at an annealing temperature of 65 °C for 15 s followed by elongation at 72 °C for 30 s. Amplicon lengths of genomic sequences were discriminated by gel electrophoresis of PCR products. Sanger sequencing of PCR amplicons was performed using Mix2seq kits at Eurofins Genomics Germany GmbH (Ebersberg, Germany).

### Whole-genome sequencing

DH lines KE0095 and KE0109, from which recombinant lines and NILs were derived, were sequenced using PacBio HiFi long reads by CNRGV, INRA Occitanie Toulouse, France (cnrgv.toulouse.inra.fr). Circular consensus sequences were *de-novo* assembled in contigs using the hifiasm assembler with default parameters (Cheng et al., 2021). Subsequently, the contigs were ordered in ALLMAPS (Tang et al., 2015) using a genetic map derived from the cross of inbred lines EP1xPH207 as reference (Haberer et al., 2020).

### Comparative genomic analyses

Genomic positions of KASP markers in the QTL region on chromosome 10 of KE0095 and KE0109 were obtained by mapping their probe sequences against KE0095 and KE0109 PacBio HiFi genomes using bwa-mem (Li and Durbin, 2009). Genomic positions of B73v5 genes (Hufford et al., 2021) in KE0095 and KE0109 were identified using BLAST+ 2.9.0 (Camacho et al., 2009).

Pairwise sequence alignment of chromosome 10 between KE0095 and KE0109 was conducted using nucmer with a minimum length of a cluster set to 156 and a maximum gap between adjacent matches in a cluster set to 1000, and subsequently filtered using delta-filter with settings of minimum identity of 95% and minimum alignment length of 1000 (Kurtz et al., 2004).

To compare the amino acid sequences of candidate genes in the QTL region on chromosome 10 between KE0095 and KE0109, coding sequences of the candidate genes were extracted from the respective genome assembly. Coding sequences were translated to amino acids and aligned using Clustal Omega (Sievers and Higgins, 2018).

LHCII components of Arabidopsis were identified in Araport 11 (Table S7; Cheng et al., 2017) and searched against B73v5 proteins (Hufford et al., 2021) via BLAST+ 2.9.0 (Camacho et al., 2009). The search result was filtered for minimum sequence identity of 70% and minimum query coverage of 75%. For each query, the search result with the highest bit score was selected and reported (Table S7).

### Haplotype distribution in elite lines

Data for 136 diverse Dent and Flint inbred lines genotyped with the 600k Array were obtained from Unterseer et al. (2016) and merged with the 600k SNP data from the Kemater DH lines, retaining only SNPs present in both datasets. The 15 SNPs closest to *lhcb6* in B73v5 (8 upstream; 7 downstream), including the most significant SNP from the GWAS analysis (AX- 90599221; lead SNP), were concatenated to define haplotype alleles. These haplotype alleles were then filtered to keep only those occurring at least three times in the combined dataset.

### Statistical analyses

All statistical analyses were performed in R (R Core Team, 2013). The statistical model for estimating genotype and genotype by environment variance components of F_2:3_ recombinant lines derived from the cross of the DH lines KE0095 and KE0109 in field experiments 2024 was:

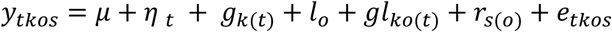

where 𝑦_𝑡𝑘𝑜𝑠_ are the phenotypic observations; 𝜇 is the overall mean; 𝜂 _𝑡_ is the fixed effect of the group 𝑡 (𝑡 =1 for checks, 𝑡 =2 for F_2:3_ recombinant lines); 𝑔_𝑘(𝑡)_ is the random effect of genotype 𝑘 nested within group 𝑡; 𝑙_𝑜_ is the random effect of environment 𝑜; 𝑔𝑙_𝑘𝑜(𝑡)_ is the random interaction effect for genotype 𝑘 and environment 𝑜 nested within group 𝑡; 𝑟_𝑠(𝑜)_ is the random effect of the block 𝑠 nested within environment 𝑜 and 𝑒_𝑡𝑘𝑜𝑠_ is the residual error. Genotypic and genotype by environment variance components were modeled individually for recombinant lines and for checks. Restricted maximum-likelihood estimation implemented in ‘ASRemlR’ (v4.1.0) was used for estimating variance components and their standard errors. Trait heritability (h^2^) and associated confidence intervals were calculated on an entry mean basis applying established procedures (Knapp et al., 1985, Hallauer et al., 2010).

Adjusted entry means for the F_2:3_ recombinant lines, the subset of 27 Kemater lines, and respective checks were calculated using the same model but dropping the group effect (𝜂 _𝑡_) and treating genotype as a fixed effect. Adjusted means for KE0095, KE0109 and four NILs in the growth chamber experiments were calculated using a statistical model with genotype as fixed effect and block as random effect.

To evaluate the association between marker alleles and phenotypic data in the F_2:3_ recombinant lines, significance of marker effects was assessed for 14 markers (Table S1) based on adjusted means with incremental Wald tests implemented in ‘ASRemlR’ (v4.1.0) (Kenward and Roger, 1997). Marker alleles were coded as 0 and 2 and marker effects were considered significant at a FDR of 1%. Phenotypic variance explained (PVE) by the markers was calculated as the coefficient of determination (R^2^) of the regression of F_v_/F_m_ adjusted means on marker alleles. Genetic variance explained (GVE) was calculated as PVE divided by the heritability estimate of F_v_/F_m_ (Schön et al., 2004). To assess the effect of the *lhcb6-B* allele on Fv/Fm and early plant height in the subset of 27 Kemater lines, a two-way analysis of variance (ANOVA) was conducted with the genes *lhcb6* and *ndhm1* (Urzinger et al., 2025), and their interaction as factors. Significance for the traits F_v_, F_m_, F_v_/F_m_, NPQ, fresh and dry biomass and half rise time from F_0_ to F_j_ in comparisons of KE0095, KE0109 and four NILs in the growth chamber experiments was tested with Student’s t-tests with Benjamini-Hochberg correction to control for multiple testing.

To assess the significance of differential protein accumulation in the 27 Kemater lines, an ANOVA was conducted for each protein separately. To avoid false positive associations, proteins were considered differentially accumulated at a FDR of 20%.

## Acknowledgements

We thank Brigitte Neuhauser, Iris Prücklmaier, Sylwia Schepella, Stefan Schwertfirm and Margot Siebler for technical assistance. We thank the Plant Technology Center (Technical University of Munich, Germany) for providing infrastructure and technical support during greenhouse, growth chamber and field experiments. We would like to acknowledge the support of William Marande and Caroline Callot from CNRGV, INRAE (http://cnrgv.toulouse.inrae.fr/) for providing assistance in PacBio long read sequencing and GENTYANE platform of Clermont-Ferrand INRAE Center (http://gentyane.clermont.inra.fr/) for providing access to PacBio sequencer.

## Funding

This study was funded by the Federal Ministry of Education and Research (BMBF, Germany) within the scope of the funding initiative “Plant Breeding Research for the Bioeconomy” (Funding ID: 031B0195, 031B0882 and 031B1301) as part of the project MAZE (www.europeanmaize.net). A Ph.D. fellowship for L.W. was funded by the Elitenetzwerk Bayern in the scope of the project “The Proteomes That Feed the World”. KWS SAAT SE & Co. KGaA funded Ph.D. fellowships for S.U. and M.M.

## Conflict of interest statement

The authors declare that they have no conflicts of interest related to the research, authorship, or publication of this manuscript.

## Author contributions

C.-C.S., V.A., M.O., P.W., L.W. and S.U. designed the research and developed ideas; L.W., S.U., V.A. and B.O. designed and performed phenotyping experiments and analyzed the data; S.R., L.W. and S.U. analyzed whole genome sequencing data; S.B. acquired and analyzed proteomics data. L.W. and S.U. conducted candidate gene analysis; L.W., S.U., M.M., M.O., T.P., D.S. and C.U. developed the plant material; L.W., S.U, V.A., P.W. and C.-C.S. wrote the manuscript; all authors read and approved the final manuscript; C.-C.S. agrees to serve as the author responsible for contact and to ensure communication.

---

**Figure S1.**
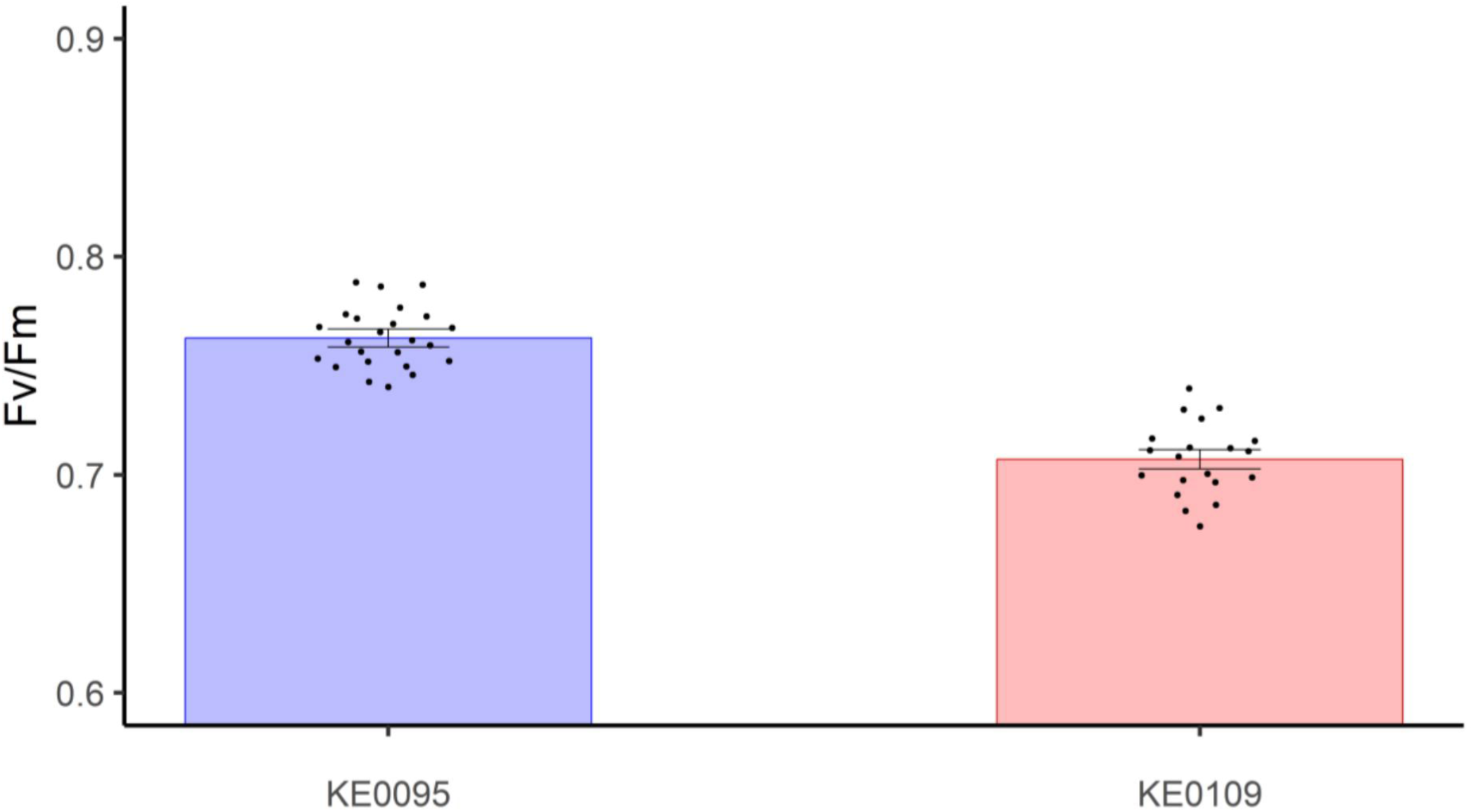
Significant difference in F_v_/F_m_ between the parental lines (KE0095 and KE0109) of the bi-parental population evaluated in a growth chamber experiment. Bars represent adjusted means ± standard errors and dots represent individual plant observations.

**Figure S2.**
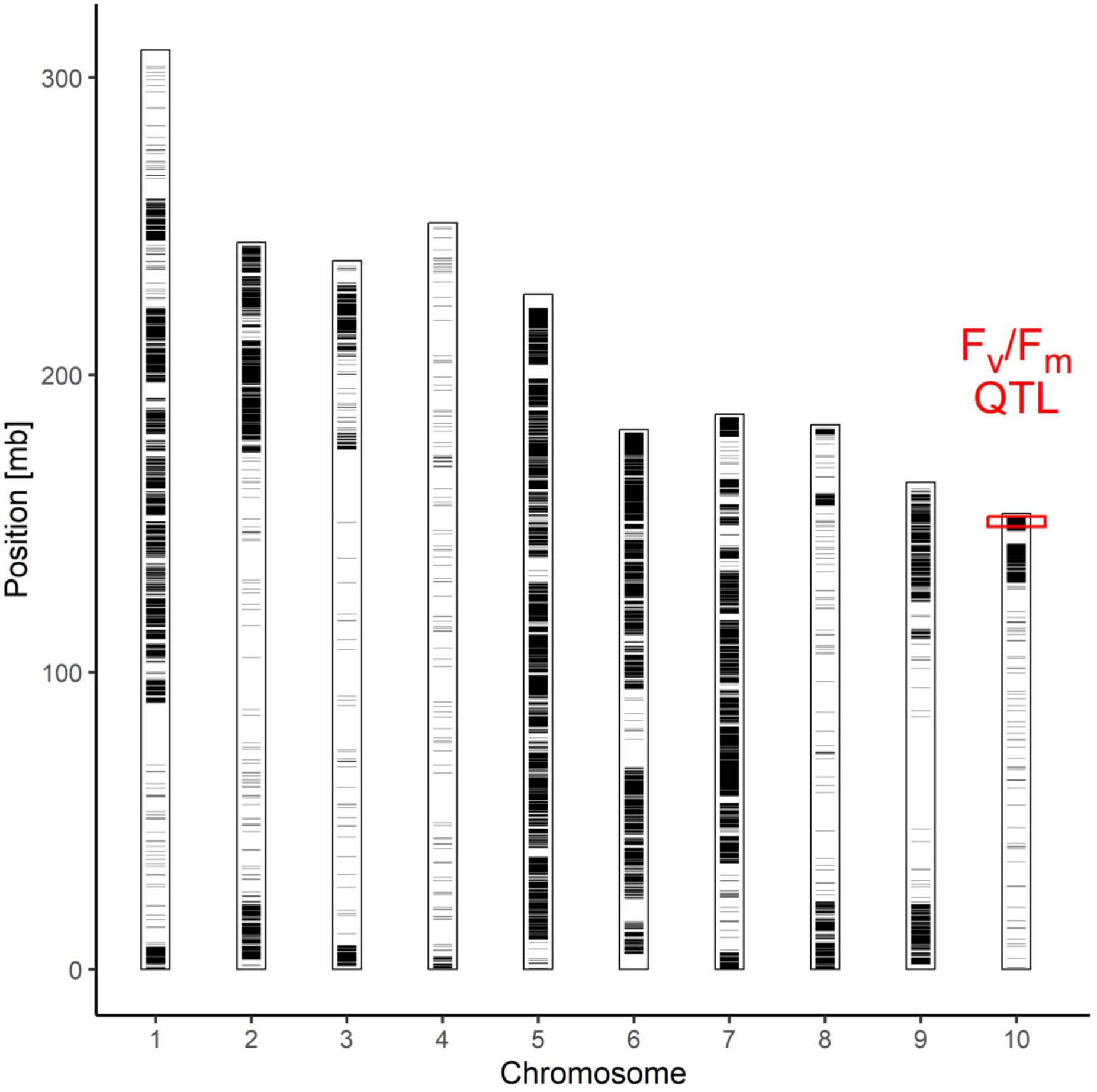
Map position of 74795 SNPs differentiating DH lines KE0095 and KE0109, the parental lines of the bi-parental population used for fine-mapping. The QTL for F_v_/F_m_ on chromosome 10 is highlighted with a red box.

**Figure S3.**
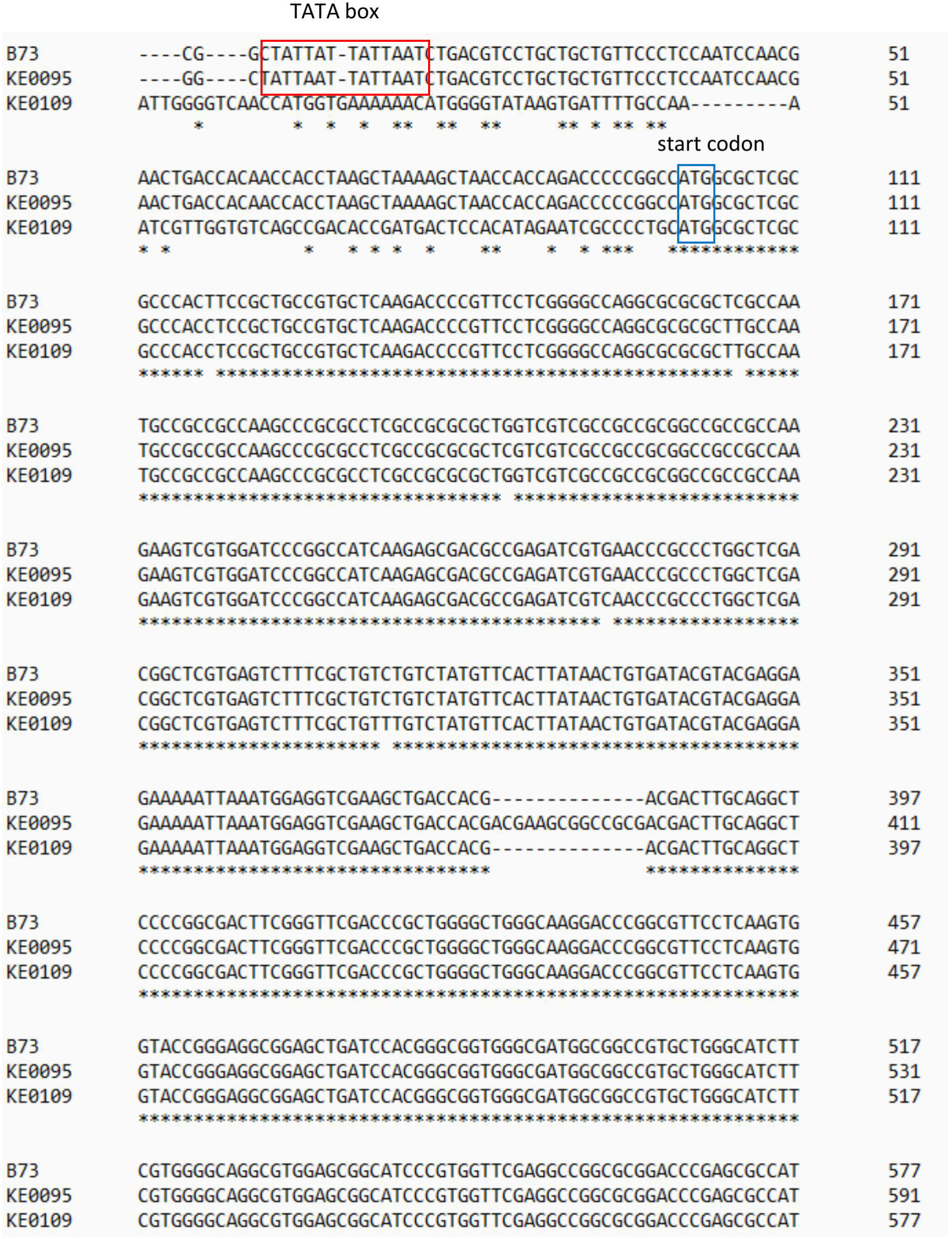
Comparison of *lhcb6* genomic sequences from B73, KE0095 and KE0109. The B73 genomic DNA sequence was obtained from MaizeGDB (B73v5) for the gene model *Zm00001eb433540* (Hufford et al., 2021). The corresponding genomic sequences for KE0095 and KE0109 were identified via BLAST using the coding sequence of *Zm00001eb433540,* including 100 bp upstream of the start codon. In the alignment, the putative TATA box is highlighted in red, and the start codon is marked in blue.

**Figure S4.**
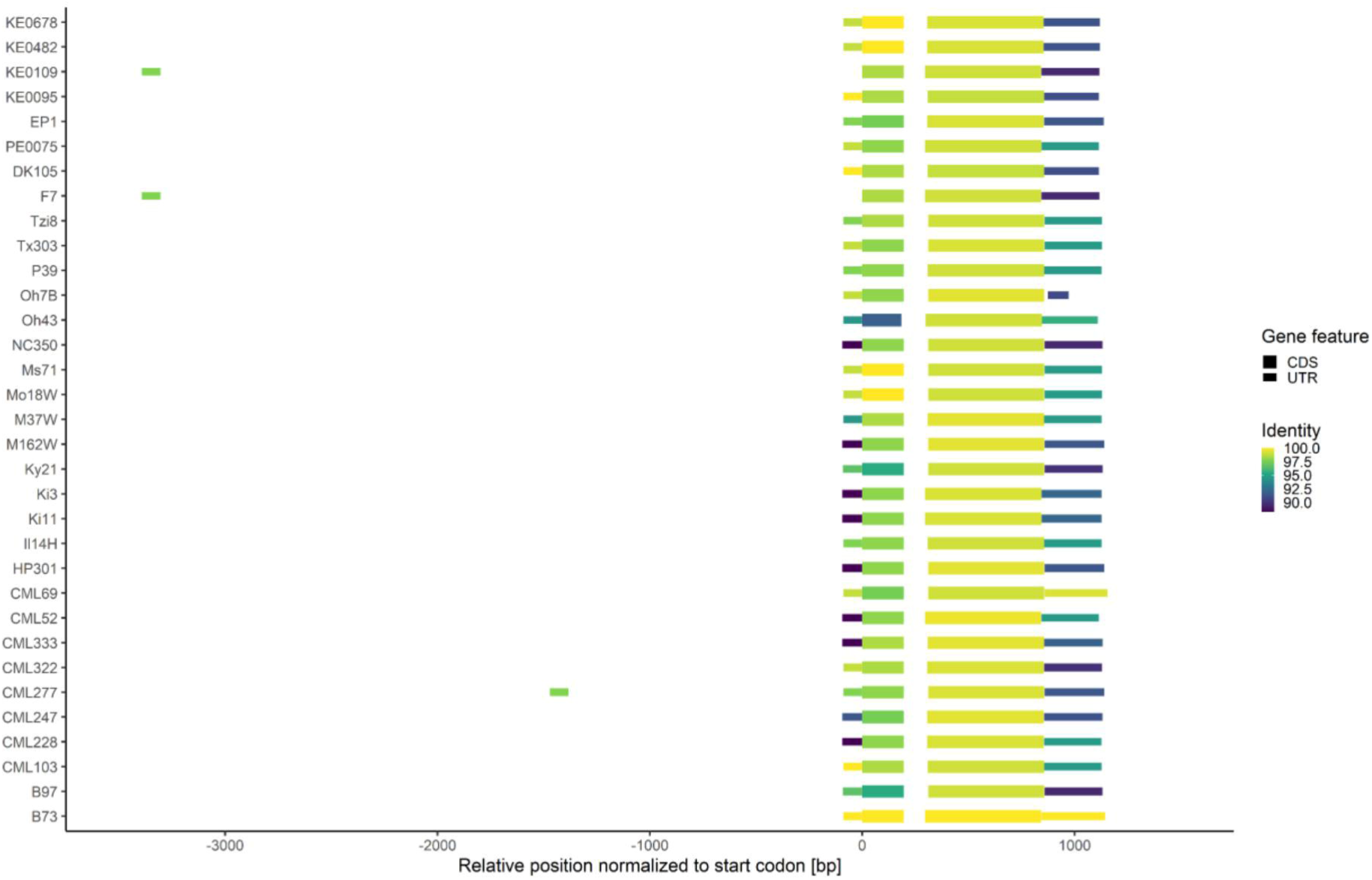
Structural comparison of *lhcb6* gene models across diverse maize genome assemblies. Genomic sequences corresponding to the 3’ UTR, 5’ UTR, and coding sequences (CDS) of *lhcb6* from B73v5 *(Zm00001eb433540; Hufford et al., 2021)* were used as queries in BLAST searches against each genome assembly. For each gene feature, only the top BLAST hit was retained, and its position was normalized relative to the start codon. Colors indicate the level of identity with the respective B73 gene feature.

**Figure S5.**
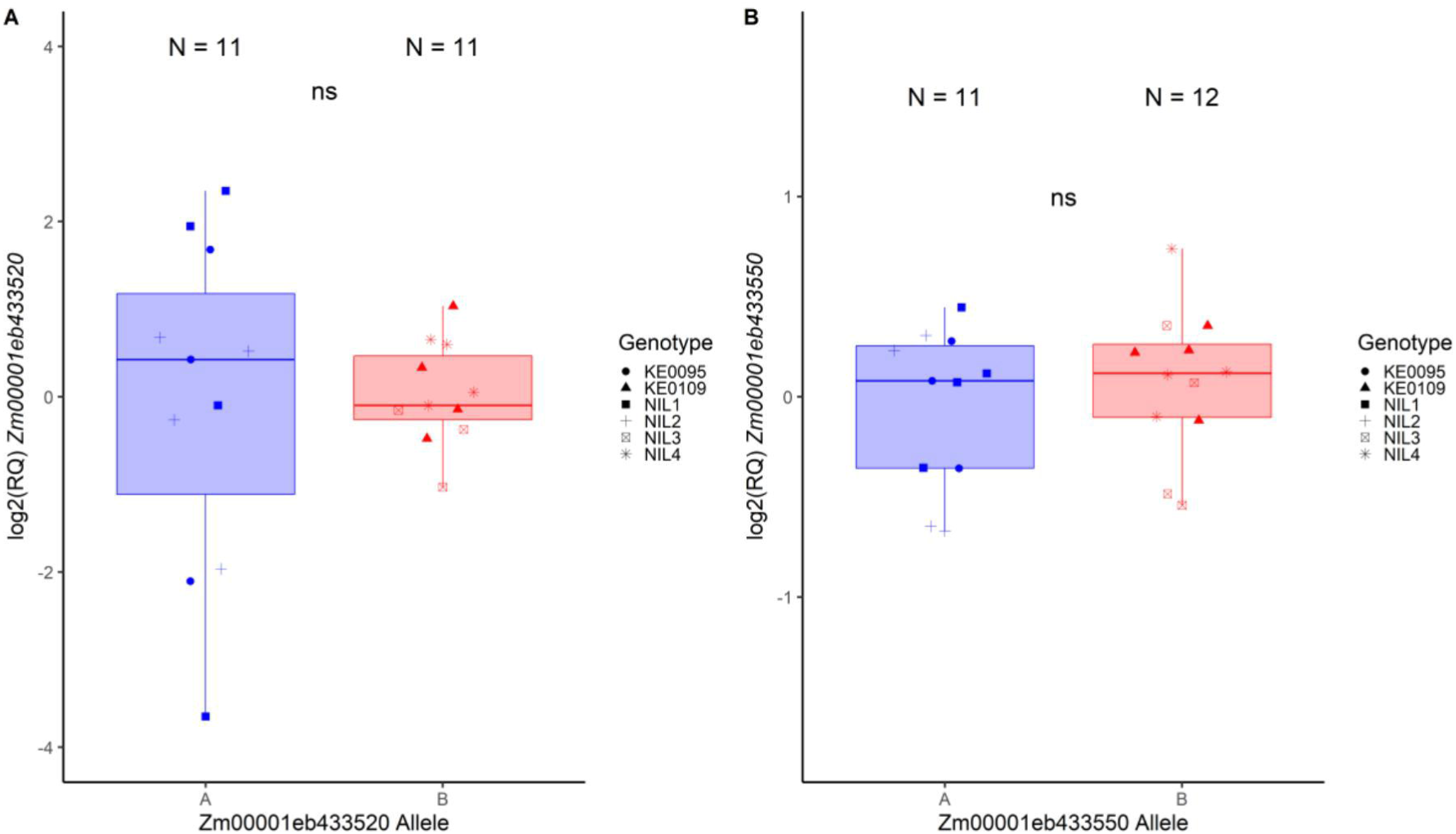
Transcript levels of candidate genes *Zm00001eb433520* and *Zm00001eb433550.* **A,B)** Log2-transformed relative transcript levels of *Zm00001eb433520* **(A)** and *Zm00001eb433550* **(B)** in near isogenic lines (NILs) with contrasting *lhcb6* alleles and of parental lines KE0095 and KE0109. Boxplots with median as center line, box limits indicate upper and lower quartiles and whiskers l.5× interquartile range. Statistical differences were assessed by analyses of variance (ANOVA). ns: not significant.

**Figure S6.**
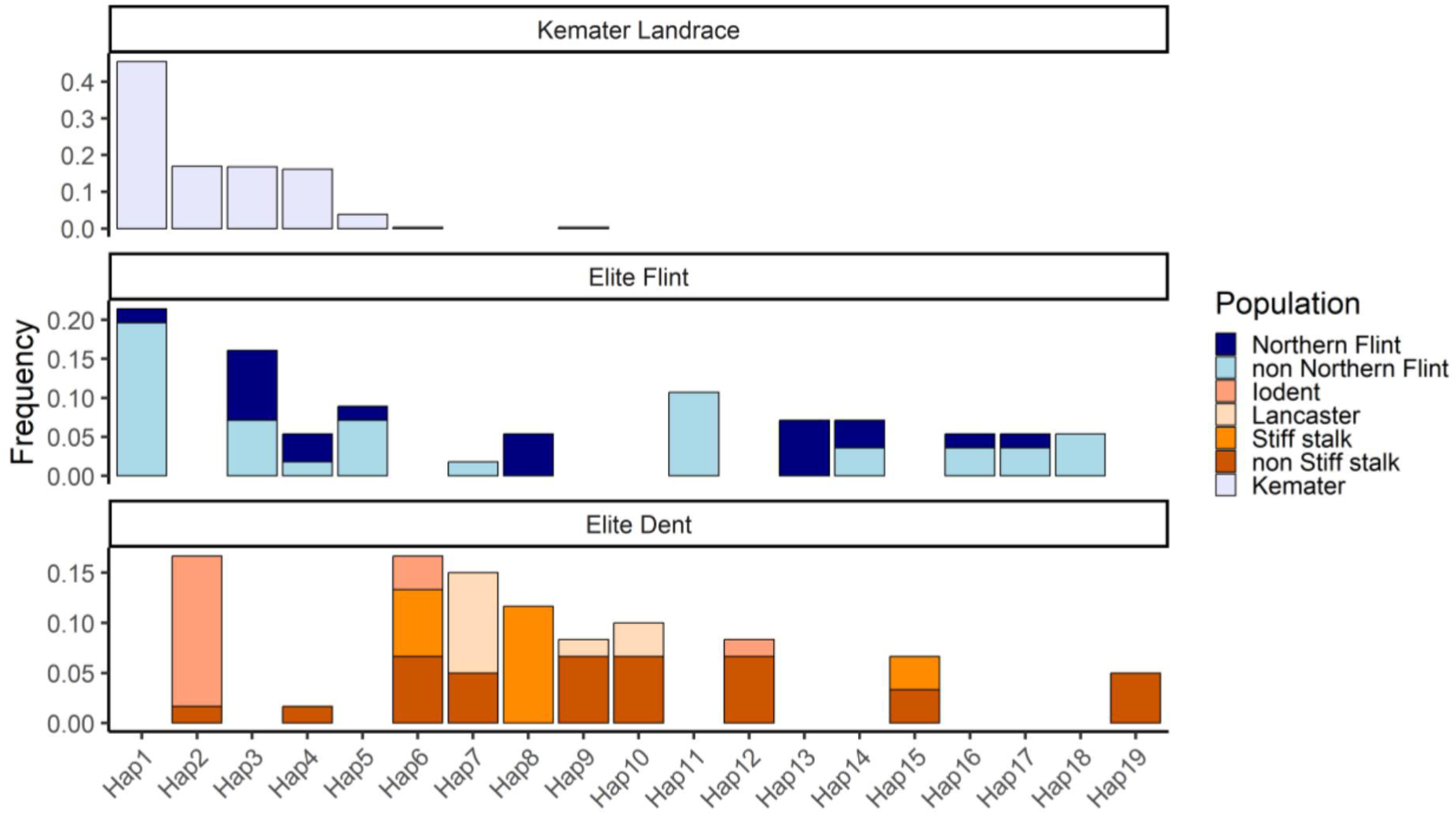
Diversity of *lhcb6* haplotypes in inbred maize lines and the landrace Kemater. Haplotypes were defined by concatenating the 15 SNPs closest to *lhcb6* from the 600k Axiom™ Maize Genotyping Array (Unterseer et al., 2014) and filtered to include only those occurring at least three times. Genotyping data and population assignments for 136 inbred lines were obtained from Unterseer *et al*. (2016). The Kemater landrace and heterotic groups of elite lines are indicated by colors.

**Figure S7.**
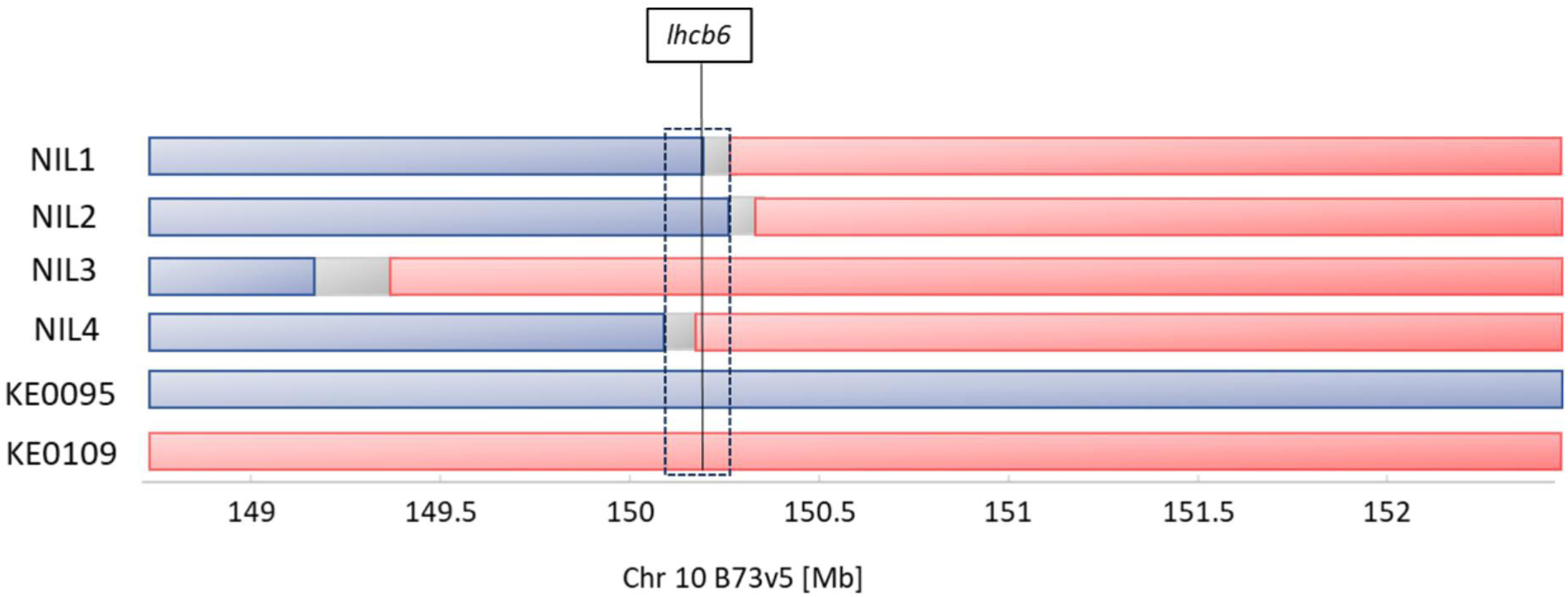
Genetic composition of near isogenic lines (NILs) and doubled-haploid (DH) parental lines used for physiological analysis in controlled conditions. Genomic regions carrying the allele of the recurrent parent (KE0095) are shown in blue, while those with the donor parent (KE0109) allele are shown in red. A 154 kb region distinguishing NIL 1 and NIL 2 from NIL 3 from NIL 4 and containing 13 gene models in the B73v5 reference genome is highlighted with a dashed box. The position of the gene *lhcb6 (light-harvesting chlorophyll a/b binding protein* 6, Zm00001eb433540) is marked with a vertical line.

**Figure S8.**
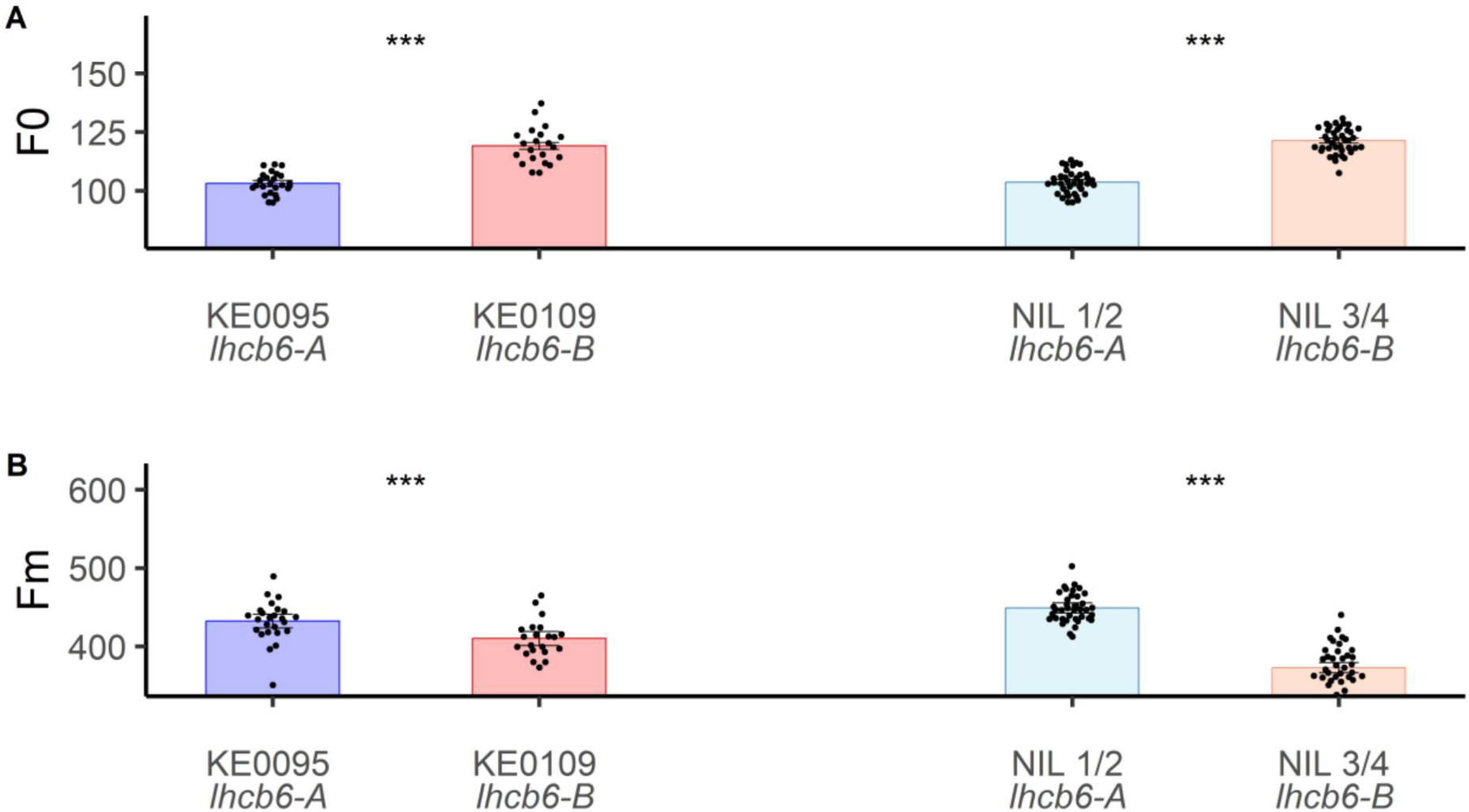
Minimal (F_0_) and maximal (F_m_) fluorescence of near isogenic lines (NILs) with contrasting *lhcb6* alleles and their parents in a growth chamber experiment. **A**) F_0_ of dark-adapted leaves. **B)** F_m_ after applying a saturating light flash to the dark-adapted leaves. Measurements were taken at growth stages V5–V6. Significant differences based on student’s t-tests are indicated by stars.** P < 0.01; ***P < 0.001;. Bars represent adjusted means ± standard error and dots represent individual plant observations.

**Table S1.** Genomic positions of KASP markers used to genotype F_2:3_ recombinant lines and near isogenic lines (NILs) from the cross of KE0095 and KE0109. Positions were determined by mapping the probes of the publicly available Axiom™ Maize 600k Genotyping Array (Unterseer *et al*., 2014) to the respective genome assembly using the bwa-mem alignment tool.

| Marker | Chromosome | Position<br>B73v5 | Position<br>KE0095 | Position<br>KE0109 | Experiment |
| --- | --- | --- | --- | --- | --- |
| AX-90560736 | 10 | 143,112,717 | 143,319,871 | 142,274,063 | NIL |
| AX-90598666 | 10 | 149,378,445 | 148,740,234 | 148,284,863 | F<sub>2:3</sub>, NIL |
| AX-91370210 | 10 | 149,463,669 | 148,872,430 | 148,442,955 | F<sub>2:3</sub>, NIL |
| AX-90611044 | 10 | 149,568,533 | 148,960,964 | 148,502,800 | F<sub>2:3</sub>, NIL |
| AX-90560838 | 10 | 149,660,591 | 149,098,879 | 148,675,336 | F<sub>2:3</sub>, NIL |
| AX-91365760 | 10 | 149,850,831 | 149,301,235 | 148,822,092 | F<sub>2:3</sub>, NIL |
| AX-91196016 | 10 | 149,960,879 | 149,413,540 | 148,934,912 | F<sub>2:3</sub>, NIL |
| AX-91369070 | 10 | 150,109,398 | 149,571,593 | 149,028,281 | F<sub>2:3</sub>, NIL |
| AX-90599221 | 10 | 150,176,073 | 149,731,916 | 149,097,971 | F<sub>2:3</sub>, NIL |
| AX-91196092 | 10 | 150,262,989 | 149,805,780 | 149,172,008 | F<sub>2:3</sub>, NIL |
| AX-90564314 | 10 | 150,367,233 | 149,893,811 | 149,194,873 | F<sub>2:3</sub>, NIL |
| AX-91196161 | 10 | 150,507,597 | 150,023,962 | 149,341,009 | F<sub>2:3</sub>, NIL |
| AX-90560860 | 10 | 150,712,585 | 150,262,052 | 149,463,226 | F<sub>2:3</sub>, NIL |
| AX-91196242 | 10 | 150,804,175 | 150,371,931 | 149,580,278 | F<sub>2:3</sub>, NIL |
| AX-90531911 | 10 | 150,914,661 | 151,359,910 | 150,249,567 | F<sub>2:3</sub>, NIL |
| AX-91196351 | 10 | 151,042,161 | 150,667,619 | 149,852,738 | NIL |
| AX-90577486 | 10 | 151,152,169 | 151,359,910 | 150,249,567 | NIL |
| AX-91834732 | 10 | 151,308,012 | 151,558,978 | 150,439,327 | NIL |
| AX-91365786 | 10 | 151,404,551 | 151,616,333 | 150,487,914 | NIL |
| AX-91196558 | 10 | 151,738,373 | 151,883,122 | 150,922,091 | NIL |
| AX-91834854 | 10 | 151,941,918 | 152,085,768 | 151,062,065 | NIL |
| AX-91834882 | 10 | 152,116,439 | 152,337,249 | 151,249,867 | NIL |
| AX-91365808 | 10 | 152,307,284 | 152,503,216 | 151,453,405 | NIL |
| AX-91196698 | 10 | 152,357,073 | 152,552,940 | 151,512,733 | NIL |

**Table S2.** Candidate genes in the 154 kb fine-mapped region for F_v_/F_m_ on chromosome 10. The B73v5 identifiers (Hufford *et al*., 2021) of candidate genes in the fine-mapped region for F_v_/F_m_, along with their annotations from NCBI, are listed. Two of the candidate genes were identified in the leaf proteomes of the parents of the bi-parental mapping population (KE0095, KE0109) and in a set of 27 Kemater lines differing at their *lhcb6* allele. Differences in protein accumulation were tested using two-sided t-tests. ns: no significant differences in protein levels between groups. ***P < 0.001; nd: not detected.

| Gene | Annotation | Proteomics<br>KE0095 - KE0109 | Proteomics<br>27 Kemater |
| --- | --- | --- | --- |
| Zm00001eb433490 | Putative homeodomain-like transcription factor superfamily protein | nd | nd |
| Zm00001eb433500 | crr2 - Cytokinin response regulator2 | nd | nd |
| Zm00001eb433510 | Early nodulin-like protein1 | nd | nd |
| Zm00001eb433520 | Uncharacterized protein | nd | nd |
| Zm00001eb433530 | ubiquinone biosynthesis protein COQ4 homolog mitochondrial | nd | nd |
| Zm00001eb433540 | lhcb6 - light harvesting chlorophyll a/b binding protein6 | *** | *** |
| Zm00001eb433550 | pat38 - protein S-acyltransferase38 | nd | nd |
| Zm00001eb433560 | Uncharacterized protein | nd | nd |
| Zm00001eb433570 | Thioredoxin domain-containing protein | nd | nd |
| Zm00001eb433590 | Uncharacterized protein | nd | nd |
| Zm00001eb433600 | FAD-binding PCMH-type domain-containing protein | nd | nd |
| Zm00001eb433610 | chlh1 - Mg chelatase subunit H 1 | ns | ns |
| Zm00001eb433620 | ysl10 - yellow stripe-like transporter10 | nd | nd |

**Table S3.**
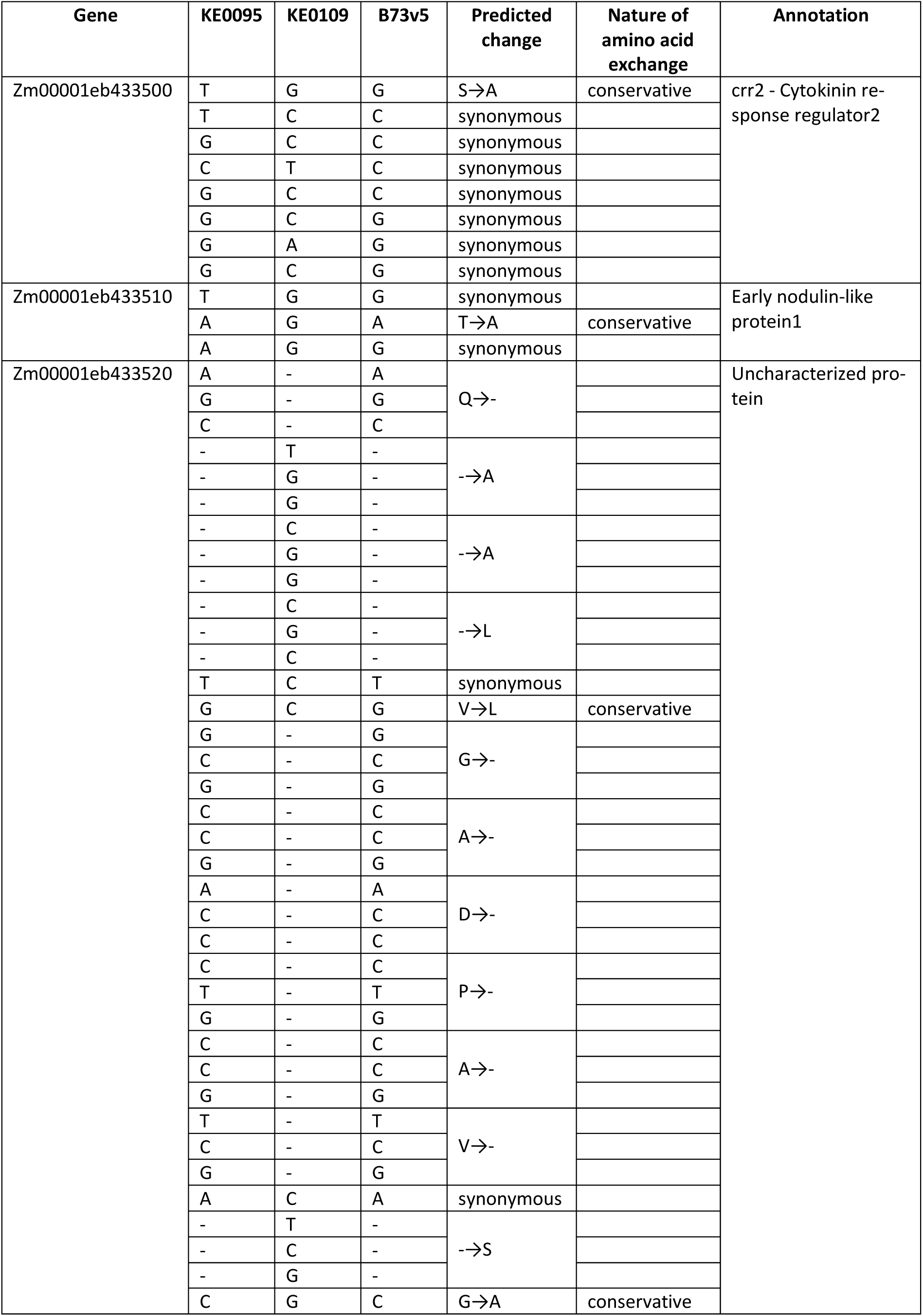
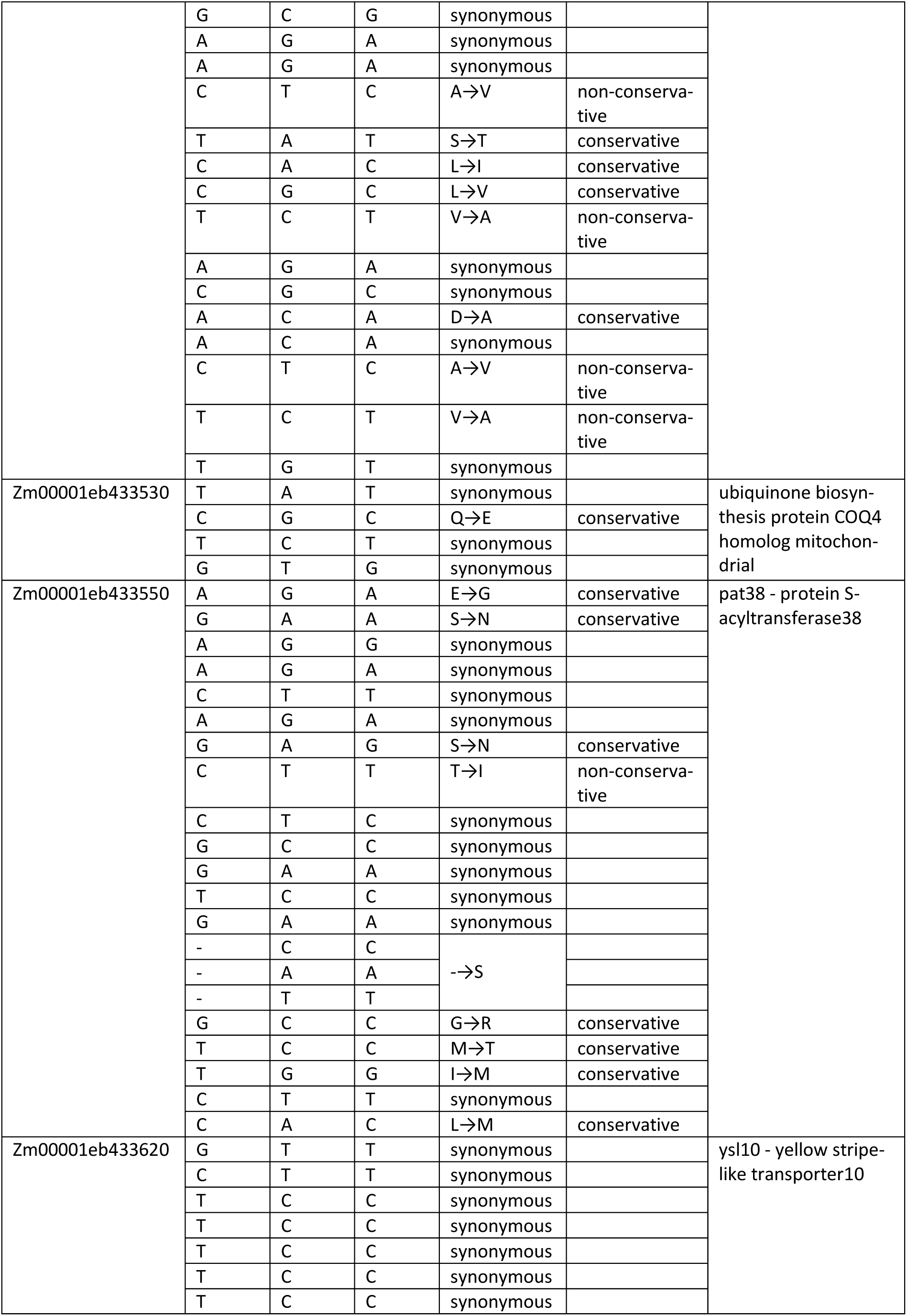
Polymorphisms between KE0095 and KE0109 in genes within the 154 kb fine-mapped region for F_v_/F_m_ on chromosome 10. For each SNP, the nucleotide of KE0095, KE0109 and B73v5 (Hufford et al., 2021) at the respective position is provided, along with the predicted amino acid exchange and whether the exchange is considered conservative.

**Table S4.** Validation set of 27 Kemater lines used for proteomic analysis and their allele at AX-90599221 (lead SNP from GWAS) and *lhcb6*. The allelic status of *lhcb6* was determined by amplifying a genomic fragment containing a hAT transposon insertion in the putative promoter of *lhcb6-B*. The *lhcb6* haplotype was determined by concatenating the 15 closest SNPs to *lhcb6* from the Axiom™ Maize 600k Genotyping Array (Unterseer et al., 2014).

| Genotype | AX-90599221 | PCR Genotyping | *Lhcb6* haplotype |
| --- | --- | --- | --- |
| KE0007 | Ade | *lhcb6-A* | Hap2 |
| KE0011 | Gua | *lhcb6-B* | Hap1 |
| KE0045 | Ade | *lhcb6-A* | Hap3 |
| KE0054 | Ade | *lhcb6-A* | Hap2 |
| KE0067 | Gua | *lhcb6-B* | Hap1 |
| KE0079 | Gua | *lhcb6-B* | Hap1 |
| KE0096 | Ade | *lhcb6-A* | Hap4 |
| KE0103 | Gua | *lhcb6-B* | Hap1 |
| KE0134 | Gua | *lhcb6-B* | Hap1 |
| KE0153 | Ade | *lhcb6-A* | Hap4 |
| KE0190 | Gua | *lhcb6-B* | Hap1 |
| KE0194 | Gua | *lhcb6-B* | Hap1 |
| KE0202 | Ade | *lhcb6-A* | Hap3 |
| KE0204 | Ade | *lhcb6-A* | Hap4 |
| KE0241 | Gua | *lhcb6-B* | Hap1 |
| KE0277 | Gua | *lhcb6-B* | Hap1 |
| KE0288 | Ade | *lhcb6-A* | Hap3 |
| KE0294 | Gua | *lhcb6-B* | Hap1 |
| KE0407 | Gua | *lhcb6-B* | Hap1 |
| KE0426 | Ade | *lhcb6-A* | Hap5 |
| KE0462 | Ade | *lhcb6-A* | Hap3 |
| KE0482 | Ade | *lhcb6-A* | Hap5 |
| KE0557 | Ade | *lhcb6-A* | Hap4 |
| KE0590 | Ade | *lhcb6-A* | Hap3 |
| KE0627 | Gua | *lhcb6-B* | Hap1 |
| KE0633 | Ade | *lhcb6-A* | Hap3 |
| KE0678 | Ade | *lhcb6-A* | Hap5 |

**Table S5.** Summary of linear model results assessing the effects of lhcb6 and ndhm1, on early growth traits in 27 Kemater lines. The factors lhcb6 and ndhm1 represent the allelic states of each gene. *lhcb6* has two levels (*lhcb6-A*, n = 15; *lhcb6-B*, n = 12), as well as *ndhm1* (*ndhm1-A1*, n = 15; *ndhm1-A2*, n = 12). Genotypic data for *ndhm1* were derived from Urzinger et al. (2025). Significance of the terms was assessed by a two-way analysis of variance (ANOVA). The table reports effect estimates, standard errors (SE), degrees of freedom (Df), sum of squares (SS), F-values, p-values, and model R².

| Phenotype | Term | Estimate | SE | Df | SS | F | p-value | R<sup>2</sup> |
| --- | --- | --- | --- | --- | --- | --- | --- | --- |
| **F<sub>v</sub>/F<sub>m</sub> (V6)** | Intercept | 0.770 | 0.003 |  |  |  |  | 0.74 |
| | *Lhcb6-B* | -0.018 | 0.006 | 1 | $2.33 \times 10^{-3}$ | 19.94 | $1.76 \times 10^{-4}$ | |
| | *Ndhm1-A2* | -0.034 | 0.006 | 1 | $5.03 \times 10^{-3}$ | 43.06 | $1.07 \times 10^{-6}$ | |
| | *Lhcb6-B:Ndhm1-A2* | 0.013 | 0.009 | 1 | $2.53 \times 10^{-4}$ | 2.17 | 0.15 | |
| | Residuals | | | 23 | $2.68 \times 10^{-3}$ | | | |
| **PH (V6)** | Intercept | 62.47 | 1.58 |  |  |  |  | 0.48 |
| | *Lhcb6-B* | -1.65 | 2.73 | 1 | 153.69 | 6.19 | $2.05 \times 10^{-2}$ | |
| | *Ndhm1-A2* | -6.48 | 2.73 | 1 | 371.40 | 14.96 | $7.80 \times 10^{-4}$ | |
|  | *Lhcb6-B:Ndhm1-A2* | -2.62 | 3.99 | 1 | 107.08 | 0.43 | 0.52 |  |
|  | Residuals |  |  | 23 | 570.82 |  |  |  |

**Table S6.** Introgressions from KE0109 in near isogenic lines (NILs) derived from the cross KE0095 x KE0109. Positions of KASP markers were determined by mapping the probes of the publicly available Axiom™ Maize 600k Genotyping Array (Unterseer et al., 2014) to the respective genome assembly using the bwa-mem alignment tool. AX-91196698 is the most distal KASP marker on chromosome 10.

| **NIL** | **Start of introgression from KE0109 between KASP markers** | **Positions [Chr 10 B73v5]** | **End of introgression from KE0109 between KASP marker and the end of chromosome 10** | **Positions [Chr 10 B73v5]** | ***lhcb6* allele** |
| --- | --- | --- | --- | --- | --- |
| **NIL 1** | AX-90599221 & AX-91196092 | 150,176,073 & 150,262,989 | AX-91196698 | 152,357,073 & 152,435,371 | *lhcb6*-A |
| **NIL 2** | AX-91196092 & AX-90564314 | 150,262,989 & 150,367,233 | AX-91196698 | 152,357,073 & 152,435,371 | *lhcb6*-A |
| **NIL 3** | AX-90525780 & AX-91385859 | 149,259,081 & 149,374,786 | AX-91196698 | 152,357,073 & 152,435,371 | *lhcb6*-B |
| **NIL 4** | AX-91369070 & AX-90599221 | 150,109,398 & 150,176,073 | AX-91196698 | 152,357,073 & 152,435,371 | *lhcb6*-B |

**Table S7.**
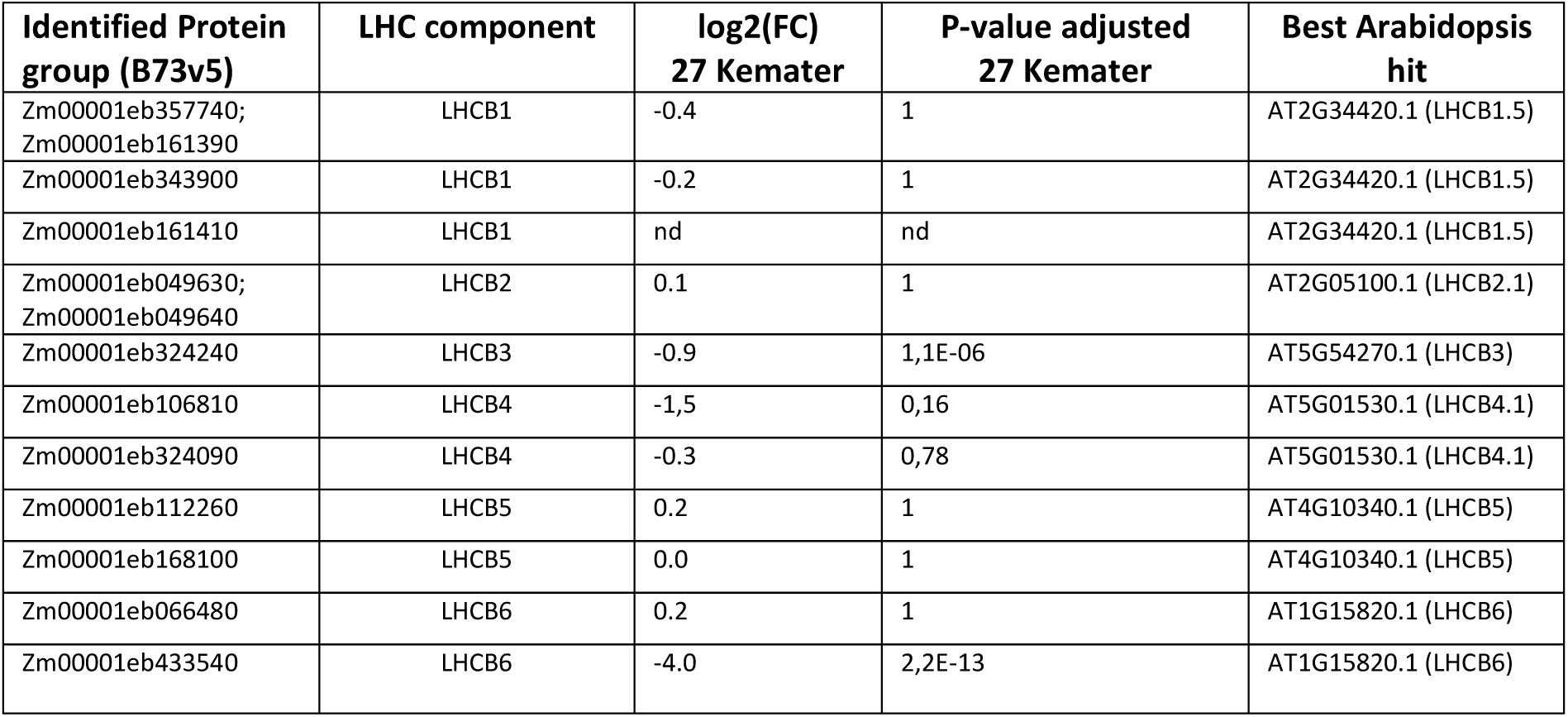
Identification of LHC II components based on homology to *Arabidopsis thaliana*. LHC II components described in *A. thaliana* were searched against proteins annotated in B73v5 (Hufford et al., 2021) using protein BLAST (Camacho *et al*., 2009). Statistical significance of differences between Kemater lines with *lhcb6-A* (N = 15) and *lhcb6-B* (N = 12) was assessed using two-sided Student’s t-tests, with resulting p-values adjusted by Bonferroni correction. Proteins that could not be distinguished by unique peptides are grouped into a single protein group. FC: fold change between *lhcb6-A* and *lhcb6-B* groups.

**Table S8.**
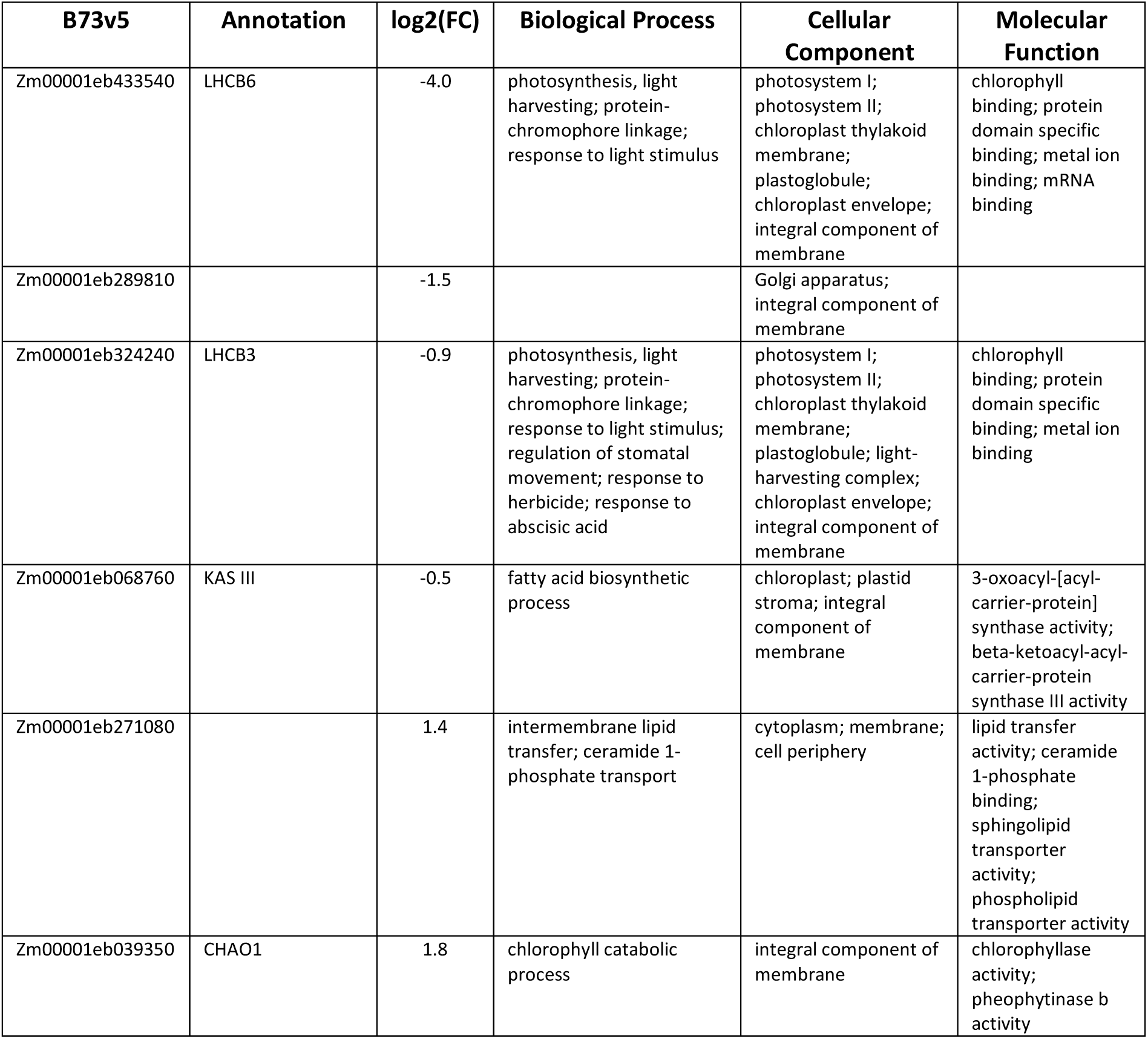
Differentially accumulated proteins in 27 Kemater lines grouped by their *lhcb6* allele. The B73v5 identifiers (Hufford et al., 2021) of differentially accumulated proteins and their associated gene ontology (GO) terms are provided. Proteins are listed in order of significance for differential accumulation. The significance of differential accumulation was determined by analyses of variance (ANOVA), with proteins selected based on a false discovery rate < 20% (Benjamini and Hochberg, 1995).

**Table S9.**
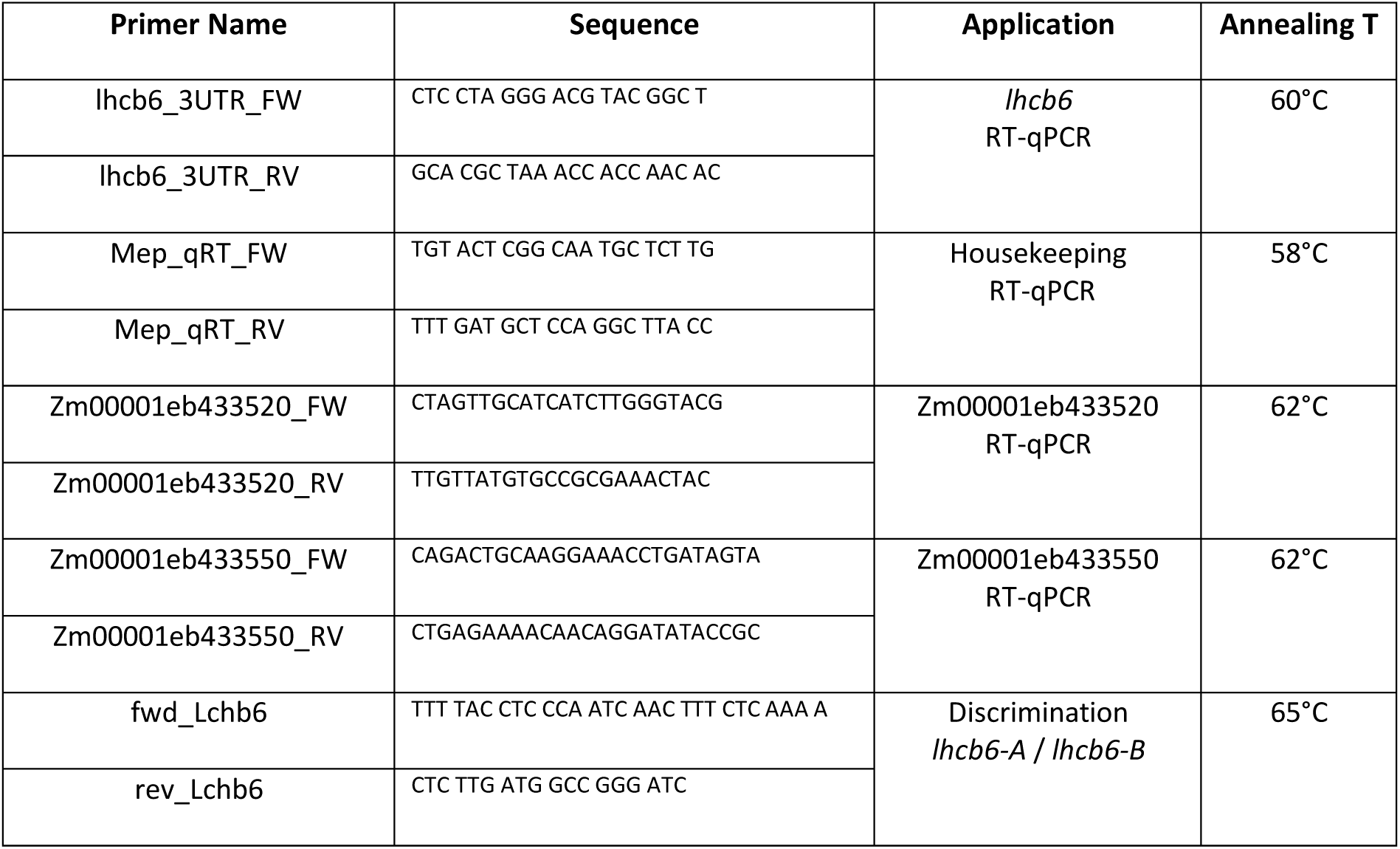
Primer used for RT-qPCR analysis and genotyping different *lhcb6* alleles. Elongation time was 30 sec for the PCRs and 60 sec for RT-qPCRs.

## Notes

### Competing Interest Statement

The authors have declared no competing interest.

